# The protocol for mesoscopic wide-field optical imaging in mice: from zero to hero

**DOI:** 10.1101/2025.07.31.667912

**Authors:** E.N. Kislukhina, N.V. Lizunova, A.M. Surin, Z.V. Bakaeva

**Author notes:** **Correspondence:** Natalia V. Lizunova. **Correspondence:** Zanda V. Bakaeva.

## Abstract

This article provides protocols that enable researchers to master mesoscopic wide-field optical brain imaging from scratch. The protocols describe surgery for wide-field cranial window creation in mice, as well as the imaging process and setup. The protocols for components of the imaging system selection and assembly, creation of a headplate for fixation, and training mice are also provided. The final section briefly outlines methods for data processing.

The described procedure can be used to visualize the dorsal cortex using wide-field optical imaging and laser-speckle contrast imaging methods. The distinguishing features of our protocol include: a wide cranial window (up to 60% of the entire cortex), skull thinning (without craniotomy), a UV-curable transparent coating (gel polish), and the ability to perform measurements in awake, behaving mice. During the surgery, a helicopter-shaped headplate with a lower surface congruent to the skull surface is mounted on the mouse’s head. This lightweight headplate allows for secure head fixation during movement eliminating the need for alignment duping data analysis. Cranial window remains sufficiently transparent for at least three months.

Wide-field optical imaging enables the recording of brain haemodynamics and energy metabolism (FAD concentration dynamics) in wild-type mice. The use of transgenic animals expressing genetically encoded sensors allows for the measurement of ions concentrations (e.g., Ca^2+^-dynamics) and other compounds (e.g., glutamate). This article describes the simultaneous measurement of changes in oxy-, deoxy-, and total haemoglobin concentrations in combination with various intracellular parameters: Δ[FAD], Δ[Ca^2+^], or ΔpH with Δ[Cl^−^].

## Introduction

*In vivo* imaging techniques are rapidly advancing in biological research, offering significant advantages over traditional histological and biochemical methods. These techniques allow for chronic studies within the same animal, reducing the number of subjects required and enhancing data quality. Notably, *in vivo* brain imaging can now be performed in awake animals, eliminating anaesthesia-related confounds (Khiroug & Verkhratsky, 2021).

Among the neuroimaging techniques, the following can be highlighted: traditional methods, used in both experimental and clinical contexts: structural and functional MRI, computed tomography, electroencephalography, electrocorticography, implanted electrodes, and ultrasound (Doppler) imaging. However, optical methods have gained prominence in experimental neuroscience, including two- and three-photon microscopy, miniscopy, Raman microspectroscopy, laser speckle contrast imaging (LSCI), and wide-field optical imaging (WFOI) (Fedotova et al., 2023; Markicevic et al., 2021). A recent achievement is the application of optical techniques (functional near-infrared spectroscopy – fNIRS) in humans (Li et al., 2022). WFOI, employing a cortex-wide cranial window, enables high spatial and temporal resolution imaging but is limited by light scattering, restricting depth measurements to the cerebral cortex (Padawer-Curry et al., 2023).

Brain activity can be observed due to the optical properties of haemoglobin, FAD and artificially created biosensors. The excitation of neurons causes vasodilation and an increase in local blood flow by the mechanism of neurovascular coupling. This is accompanied by an increase in total (HbT) and oxygenated haemoglobin (HbO), and a decrease in deoxyhaemoglobin (HHb). In addition, electrical activity requires the consumption of energy substrates (e.g., reduced flavine adenine dinucleotide), which is followed by an increase in the ratio [FAD]/[FADH_2_]. Changes in FAD and haemoglobin concentrations can be measured photometrically, since haemoglobin is a pigment and absorbs light in the visible part of the spectrum (with distinct absorption spectra for HbO and HHb), and FAD (but not FADH_2_) has fluorescent properties (Chance et al., 1979; Kozberg et al., 2016). The use of genetically encoded fluorescent sensors allows researchers to expand the panel of visualised processes. Among all genetically encoded sensors, the most widely used is the Ca^2+^ sensor GCaMP (Dana et al., 2014; Y. Zhang et al., 2023), however, there are also sensors for pH, Cl^−^, glutamate, etc (Arosio et al., 2010; Day-Cooney et al., 2023; Diuba et al., 2020a, 2020b; Kotova et al., 2023; Marvin et al., 2018). An increase in the concentration of a particular ion or molecule causes an increase in fluorescence or quenching of the sensor. For such experiments, transgenic mouse lines are used, or viral transduction is carried out to express protein sensors in brain cells. All optical neuroimaging methods require the creation of a cranial window to enable optical access to the brain, which is achieved in various ways.

One advantage of wide-field optical neuroimaging is its ability to capture signals from a large area of a dorsal part of the cerebral cortex (dorsal cortex) simultaneously (Fig. 1). While signal detection through an intact skull is possible, it is affected by diploic vessel activity and significant light scattering. To overcome these issues, researchers either perform a craniotomy or thin the skull. The latter is a less invasive approach but requires specific skills to achieve optimal results.

**Figure 1.**
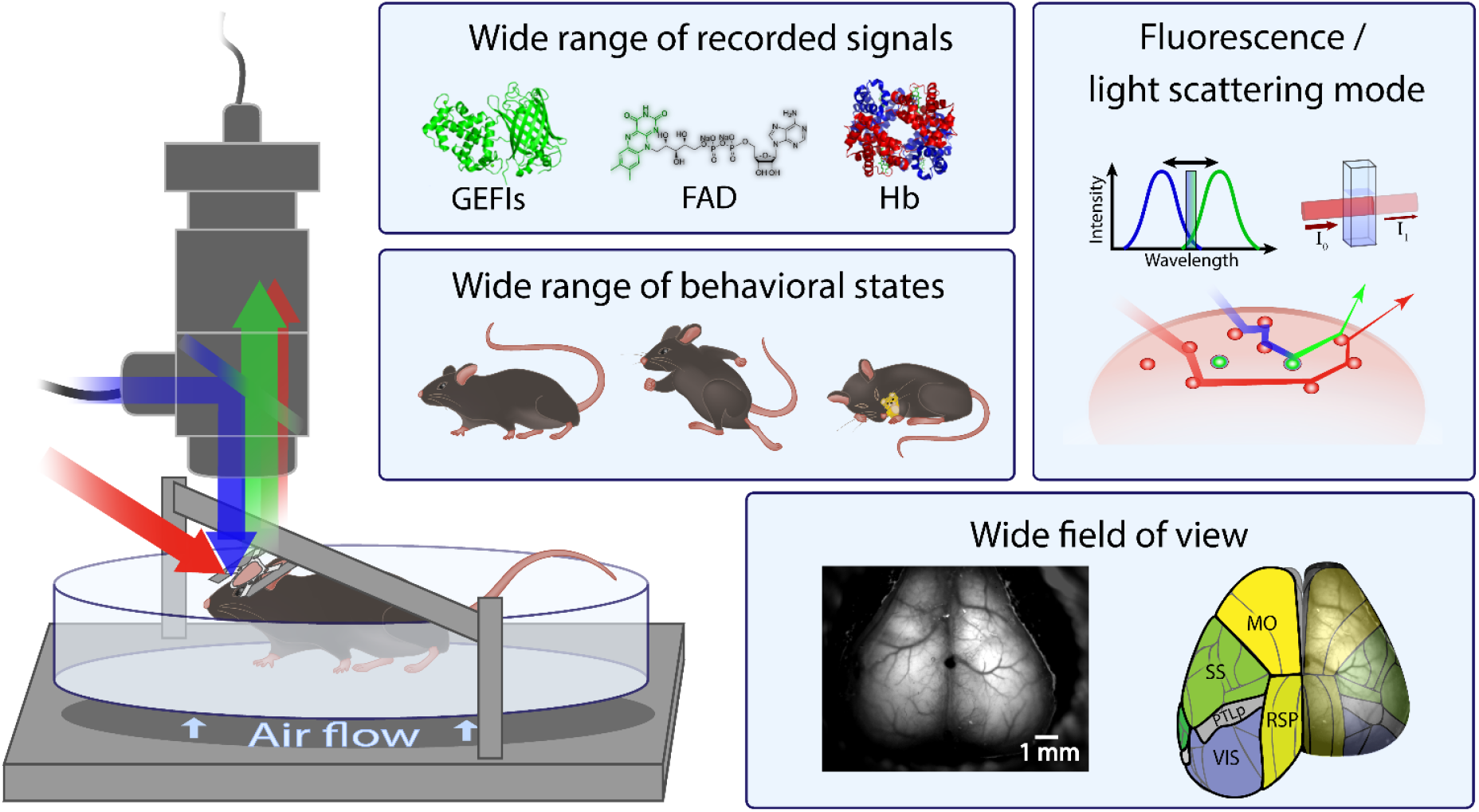
The main advantages of wide-field optical imaging. Wide-field optical imaging enables the measurement of changes in the concentrations of endogenous fluorophores (e.g., FAD) and pigments (e.g., haemoglobin, Hb), as well as signals from artificial genetically encoded fluorescent indicators (GEFIs), in both awake freely moving animals and anaesthetized ones. Measurements can be performed in fluorescence or light-scattering (reflectance, intrinsic optical signal) modes. A wide cranial window enables visualisation of all major cortical regions except the auditory cortex. MO – Somatomotor areas; PTLp – Posterior parietal association areas; RSP – retrosplenial area; SS – Somatosensory areas; VIS – visual areas, according to the Allen mouse brain common coordinate framework (Q. Wang et al., 2020).

Another key advantage is the ability to record brain activity in awake animals. This requires the development of a system that ensures stable head fixation while allowing for the simulation of natural locomotion.

Additionally, wide-field optical neuroimaging offers remarkable versatility. By integrating different LEDs (light-emitting diodes), stimulus delivery devices, and genetically encoded sensors, researchers can address a wide range of experimental questions. While commercial imaging systems are available, many laboratories prefer to assemble custom setups using various components to better suit their specific research needs.

Our protocols provide a step-by-step guide to cranial window preparation by skull thinning with a transparent protective coating, imaging system assembly, and animal training for imaging sessions. By outlining a cost-effective and adaptable approach to WFOI, we aim to make high-precision neural imaging more accessible to researchers. The protocols were extensively tested and refined. Selected results have been published in (Kislukhina et al., 2023).

## Materials and Methods

The equipment, tools, and consumables used in this study are listed in Supplementary Table S1.

### 1. Headplate design

This section describes the process of manufacturing headplates (Fig. 2) that are compatible with the Mobile Homecage® system (Neurotar, Finland). This headplate design has the following advantages: wide field of view; lightness and compactness; congruence to the surface of the skull; secure head fixation during imaging, preventing frame alignment issues during data processing.

**Figure 2.**
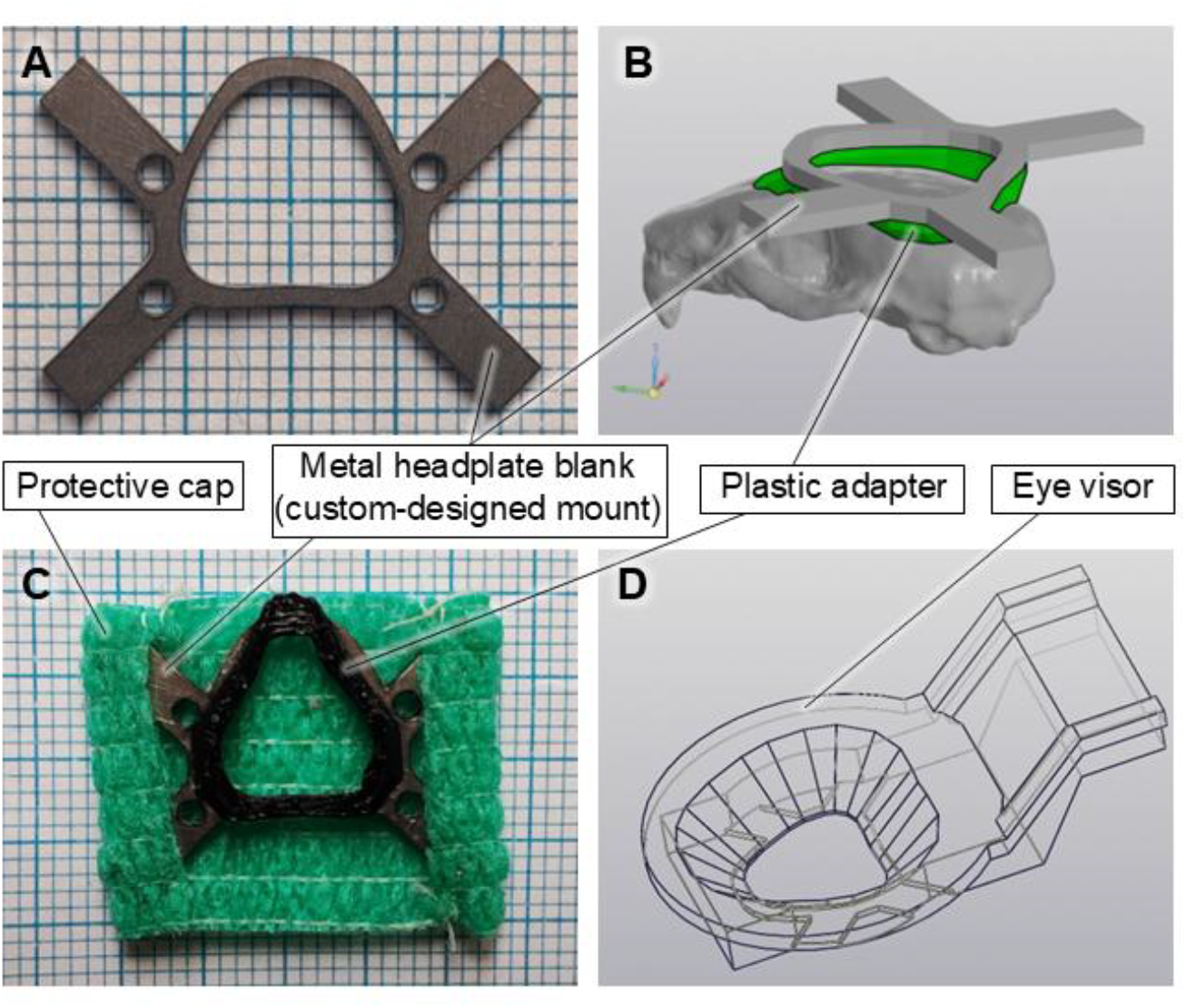
A headplate for securing an animal’s head under a microscope. A) A titanium headplate blank (custom-designed mount); B) A 3D model of an adapter that is congruent with the surface of the mouse’s skull; C) The whole headplate: metal blank with a glued plastic adapter and a removable protective cap (ventral view); D) A 3D model of a visor, which prevents eye stimulation from LEDs of the illumination system.

The protocol describes headplate manufacturing from two parts: a metal blank and a concave adapter made of plastic. At the end of the protocol, a method for manufacturing a protective cap for the brain surface, which prevents injury, is described. The Supplementary file S2 provides alternative protocols for headplate design.

#### 1.1. Manufacturing headplates

1. Order laser-cut flat reusable headplate blanks from a 1 mm thick sheet of steel or titanium according to the drawing (see Supplementary file S3). *Note: Titanium is preferred because it is lighter and does not corrode*.
2. Print the black PET-G plastic headplate adapters using a 3D printer (for the 3D model, please see Supplementary file S3). If needed, refine them with a needle file. *Note: The black colour of the plastic prevents light from penetrating through the headplate*.
3. Degrease the surfaces. Apply cyanoacrylate glue to the metal workpiece. Place the plastic headplate adapters on top to match contours of the two items. Press firmly for 10 s and allow to dry for 1–2 minutes (Fig. 2). *Note: If these headplates do not meet your requirements, please see Supplementary file S2 for alternative protocols, including the creation of custom adapters from epoxy resin*.
4. 3D-print an eye visor using black PET-G plastic (for the 3D model, please see Supplementary file S3).
5. Cut a 20 × 32 mm piece of veterinary bandage with bitter impregnation. Fold 4 mm on each side to make a 20 × 24 mm piece. Sew it up to make two pockets for the “propeller blades” of the helicopter-like headplate (Fig. 2C). Test whether it is easy to put it on and off. The cap should not touch the mouse’s eyes.

*Note: The protective cap prevents brain damage and dust ingress. The cap is essential for keeping male mice together, as they are prone to fighting. It is not strictly necessary for solitary confinement or for keeping females together*.

#### 1.2. Reuse of metal blanks

6. Remove the headplate from the mouse’s head at the end of an experiment. Place the headplates in acetonitrile for a minimum of 2 hours to dissolve dental resin and glue.
7. Remove any remaining glue and plastic with sandpaper or a needle file.

### 2. Wide-field thinned cranium surgery

This section describes a surgical procedure to create a wide cranial window (Fig. 3). The main advantages of this protocol over craniotomy include: lower invasiveness, as the cranial cavity remains intact; low risk of bleeding; and ease of coating. The main limitations include bone regrowth if the osteogenic layer is not removed completely, and the requirement for a delicate surgical technique as a neuroinflammatory response may occur. For a video protocol see https://www.youtube.com/@WIFIOPIA. Detailed techniques and material specifications are available in Supplementary table S1 and Supplementary file S2. A troubleshooting guide is in Supplementary file S4.

**Figure 3.**
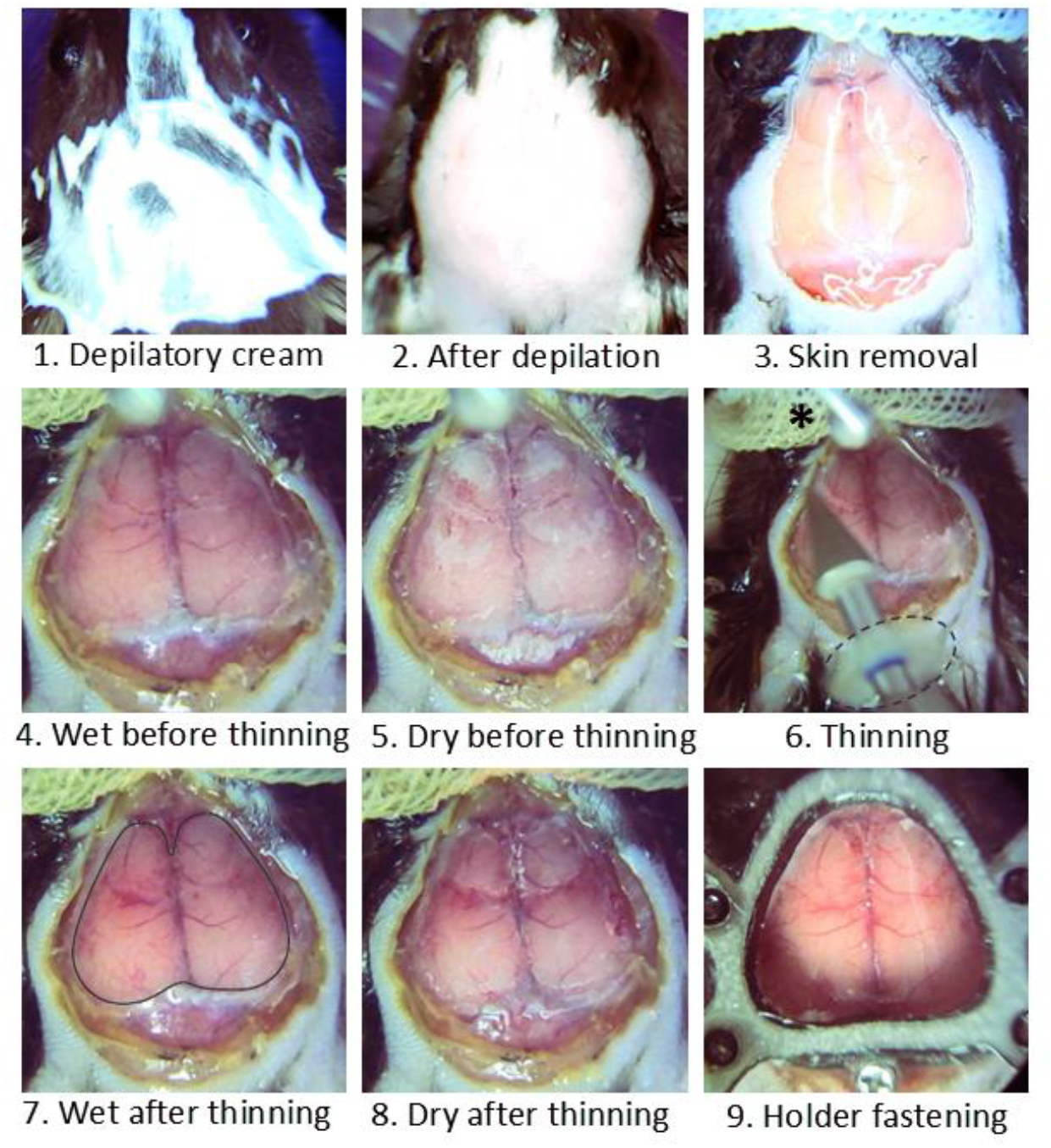
Stages of the surgery. 1. Apply depilatory cream for 1–5 minutes. 2. Remove depilatory cream with a nail stick and a damp cotton swab. 3. Sterilise surgical field; remove skin and fascia. 4. Set up continuous rinsing with saline using a peristaltic pump; use gauze for drainage. 5. Check skull thickness periodically by drying with an air flow. 6. Thin the skull using a microdrill with progressively softer milling cutters; * – tip of the saline needle and drainage gauze, cotton wool is placed underneath the head for saline absorbance; ∘ – protective “skirt” for the milling tool to prevent fluid from entering the drill. 7. The wet thinned skull has minimal differences from the intact one; ○ – thinned area. 8. Good visibility of blood vessels trough a dry scull indicates proper thinning. 9. Secure the screw into the occipital bone; fasten the holder with cyanoacrylate glue and dental resin; cover the skull with cyanoacrylate glue, base and top gel polish.

All experiments with animals should be conducted in accordance with local ethical guidelines.

#### 2.1. Preparation for surgery. Perform one day or more before surgery

1. Cover all work surfaces and equipment with plastic film, leaving the tips of ear bars, the fronts of the dental clamp, the binocular lenses, the light source, and the tip of the saline needle (see 2.1.7) uncovered.
2. Prepare protective “skirts” for milling tools by cutting approximately 7 mm plastic discs with central holes matching the cutter shaft. Secure them approximately 5 mm from the tip using gel polish. *Note: Steps 1–2 prevent fluid from entering the drill and other equipment*.
3. Prepare the following solutions aseptically (Table 1): Marbofloxacin (0.00017 mg/mL in water for injection), Ketoprofen (1.65 mg/mL in saline), and Dexamethasone (0.35 mg/mL in saline). Store at +4 °C in aliquots.
4. Prepare angled cannulae tips 18G, 45°; bend 27G needles to an L-shape for skull cleaning, glue application, bubble removal, and to a semi-C-shape for lacquer marking.
5. Create a visor from aluminium foil to protect the mouse’s eyes from UV light.
6. Cut gauze into 2 × 6 cm strips and prepare cotton swabs and cotton wool for cleaning and drainage; store small supplies in one place for convenience.
7. Route the silicone tubing from the peristaltic and air pumps to the stereotaxic frame above, attaching appropriate tips or needles for fluid delivery and airflow control.

**Table 1.** Medications.

| Reagent | Final concentration | Amount |
| --- | --- | --- |
| **Marbofloxacin 0.00017 mg/mL solution** |  |  |
| Marbobel 2 (Marbofloxacin 20 mg/mL) | 91.7% | 11 ml |
| Water for injection | 8.3% | 1 ml |
| **Ketoprofen 1.65 mg/mL solution** |  |  |
| Ketonal (Ketoprofen 50 mg/mL) | 3.3% | 0.5 ml |
| Saline (0.9% NaCl) | 96.7% | 14.5 ml |
| **Dexamethasone 0.35 mg/mL solution** |  |  |
| Dexamethasone for injection (Dexamethasone 4 mg/mL) | 8.8% | 132 µl |
| Saline (0.9% NaCl) | 91.2% | 1368 µl |

*Notes:*

– *Immediately dry and rinse equipment with distilled water if contact with saline occurs, to prevent corrosion*.
– *Marbofloxacin may precipitate in saline; use water for dilution*.

#### 2.2. Surgery

##### Preparation for surgery. Perform on the day of the operation

1. Ensure the availability of all tools, materials, and medications.
2. Disinfect the workspace, instruments, and screws (avoid heating headplates; they can be treated briefly with 70% ethanol prior to gluing).
3. Prepare materials for cleaning the depilatory cream (distilled water, nail stick, swab) and a clean cage with food and water.
4. Flush the perfusion pump tubing with 50 ml 6% peroxide, 50 ml distilled water, and 25 ml saline, unless reused consecutively.
5. Set up the perfusion pump by hanging a 200 mL saline infusion bag, and connect it to the pump tubing. Configure the flow rate to 1–2 mL/min.
6. Turn on the temperature-controlled pad (set to 37 °C).

##### Animal preparation

7. Anaesthetise the animal in the induction chamber of the isoflurane system (4% isoflurane).
8. After anaesthesia induction, quickly shave the surgical area using an electric shaver.
9. Transfer the animal to the stereotaxic frame, and reduce isoflurane to 1–2.5% to maintain stable anaesthesia.
10. Apply eye gel regularly (~every 15 min) to prevent drying.
11. Administer dexamethasone intramuscularly (0.7 mg/kg).
12. Insert and secure a lubricated rectal probe.
12. Inject 0.1 mL of 0.2% lidocaine subcutaneously under the scalp.
14. Place cotton wool under the head to absorb saline.
15. Adjust the microscope and lighting for binocular viewing.

##### Preparation of the surgical field

16. Apply depilatory cream to the scalp for 1–5 minutes (depending on its freshness), covering a slightly wider area than the planned incision; remove the cream and fur using a nail stick, then clean the skin with a damp cotton pad (Fig. 3, panels 1–2).
17. Disinfect gloves and clean the mouse’s head three times with 70% ethanol and povidone-iodine antiseptic. *Note: Avoid contact of ethanol and depilatory cream with the eyes*.
18. Confirm the absence of a pedal reflex.
19. Excise the scalp to expose the occipital, parietal, frontal, and part of the nasal bones; do not expose muscles; ensure smooth incision edges (Fig. 3, panel 3).
20. Control capillary bleeding by applying a topical haemostatic agent for 1-2 min.
21. Remove all connective tissue from the skull surface, including the edges, to ensure proper headplate fixation and prevent interference with drilling.

##### Thinning of the skull

23. Place saline-soaked gauze over the nasal bones, covering the eyes and retracting the vibrissae.
24. Ensure stable head fixation in the stereotaxic frame; reinforce the ear bar placement if needed.
25. Activate the peristaltic pump (1–2 mL/min) and adjust the saline needle to ensure that a saline drop lands rostral to the thinning site.
26. Set the microdrill speed (~25,000 rpm) and insert the medium-grit diamond bur.
27. Place the index and middle fingers of the non-dominant hand on the ear bars. Stabilize your working hand by resting the little finger and ulnar side on top of the index finger or on the heating pad, then begin skull thinning with the microdrill.
28. First, thin the frontal bones; control bleeding as needed (Fig. 3, panel 6). Maintain a tangential motion for uniform thinning.
29. Slightly thin the sutures, avoiding over-thinning near the rostral quarter of the superior sagittal sinus (Fig. 3, panel 7).
30. Stop the drill and pump; dry the skull with airflow and assess thinning. Pinkish areas indicate adequate thinning (Fig. 3, panels 4-8).
31. Resume thinning of the parietal bones with continuous movements across the sagittal suture.
32. Refine thinning of the sutures. Pay special attention to the boundaries of the frontal bones as they are prone to regrowth. Polish the surface using progressively softer cutters: medium-grit silicone-diamond polisher (blue), fine-grit (red), then extra-fine grit (grey). Leave the inner bone layer intact.
33. Continue until the skull is uniformly transparent when both wet and dry, with visible vasculature.

*Notes:*

– *Stable head fixation and stabilisation of the drilling hand reduce the risk of brain injury*.
– *Proper drainage prevents mouse aspiration and keeps optics clean*.
– *Bleeding may occur from diploic vessels, this is expected*.
– *Uneven thinning increases the risk of fracture; avoid pressure near the thinned (pinkish) areas*.
– *Avoid drying the fully thinned skull for extended periods*.

##### Application of Coating and Headplate Fixation

34. Reapply eye gel and dry the skull (Fig. 3, panel 8).
35. Apply a thin layer of cyanoacrylate glue to the thinned area using the L-shaped 27G needle. Allow it to harden.
36. Confirm head stability and insert a M1 screw into an indentation pre-drilled with a tungsten carbide drill bit in the occipital bone, without penetrating fully.
37. Use a No. scalpel blade to score nasal bones for proper headplate fixation.
38. Sterilise and test-fit the headplate, then glue it in place with gentle pressure.
39. Shield the eyes with the foil visor. Apply a layer of base coat gel polish to the thinned skull. Cure under UV light for 15s, then for 60s. Repeat with the top coat.
40. Mark Bregma with black nail polish if no longer visible. Use a semi-C-shaped needle.
41. Prepare self-curing acrylic dental resin: mix ~0.2 g of powder and 0.25 ml of liquid in 5 mL silicone mixing cup. Apply it gradually around the headplate using a syringe fitted with 18G angled cannulae tips, sealing the entire perimeter including skin edges, but avoiding contact with the cranial window or eyes (Fig. 3, panel 9).
42. After the resin sets, remove the animal from the frame. Administer marbofloxacin (8 mg/kg), ketoprofen (8.5 mg/kg), and 0.5 mL saline subcutaneously.
43. Discontinue isoflurane and place the mouse in a clean, heated single-housing cage. Monitor for at least 1 hour after the operation.
44. For the next 3 days, administer Marbofloxacin and Ketoprofen daily while monitoring recovery and inflammation.
45. Continue rehabilitation for at least 14 days (preferably up to 1 month) before experiments.

*Notes:*

– *Cyanoacrylate glue eliminates the osteoblast population*.
– *Prevent bubble formation and blood ingress in the glue, gel polish, and resin*.
– *If possible, make a flat surface with gel polish, glare may occur on a spherical surface*.
– *UV curing generates heat; use intermittent exposure to avoid thermal injury*.
– *Avoid excessive resin on the sides of the headplate to ensure proper holder mounting afterwards*.
– *Post-operative injections (antibiotic and analgesic) should be administered subcutaneously in the back area rather than into the scruff to avoid disturbing the surgical site*.

### 3. Component selection and assembly for wide-field optical imaging system

This section describes the assembly and configuration of the WFOI system. The first part provides the minimal required configuration for the measurement of changes in HHb or HbT only in an anaesthetised animal. The second part describes further modifications to enable the measurement of multiple parameters — HbO, HHb, and HbT, alongside FAD (in wild-type mice), or Ca^2+^ (in mice expressing GCaMP), or pH and Cl^−^ (in mice expressing ClopHensor) — as well as sensory stimulation in awake, behaving animals.

The system can be easily modified for laser speckle contrast imaging or adapted for different fluorescent sensors. The limitation is the short-wavelength range (up to 430 nm), as it penetrates poorly through the skull coating.

The fundamental design of the imaging system is shown in Fig. 4. Since individual components used to build imaging systems may vary across laboratories, we have focused on general principles of component selection and assembly, rather than the specific details of our setup.

**Figure 4.**
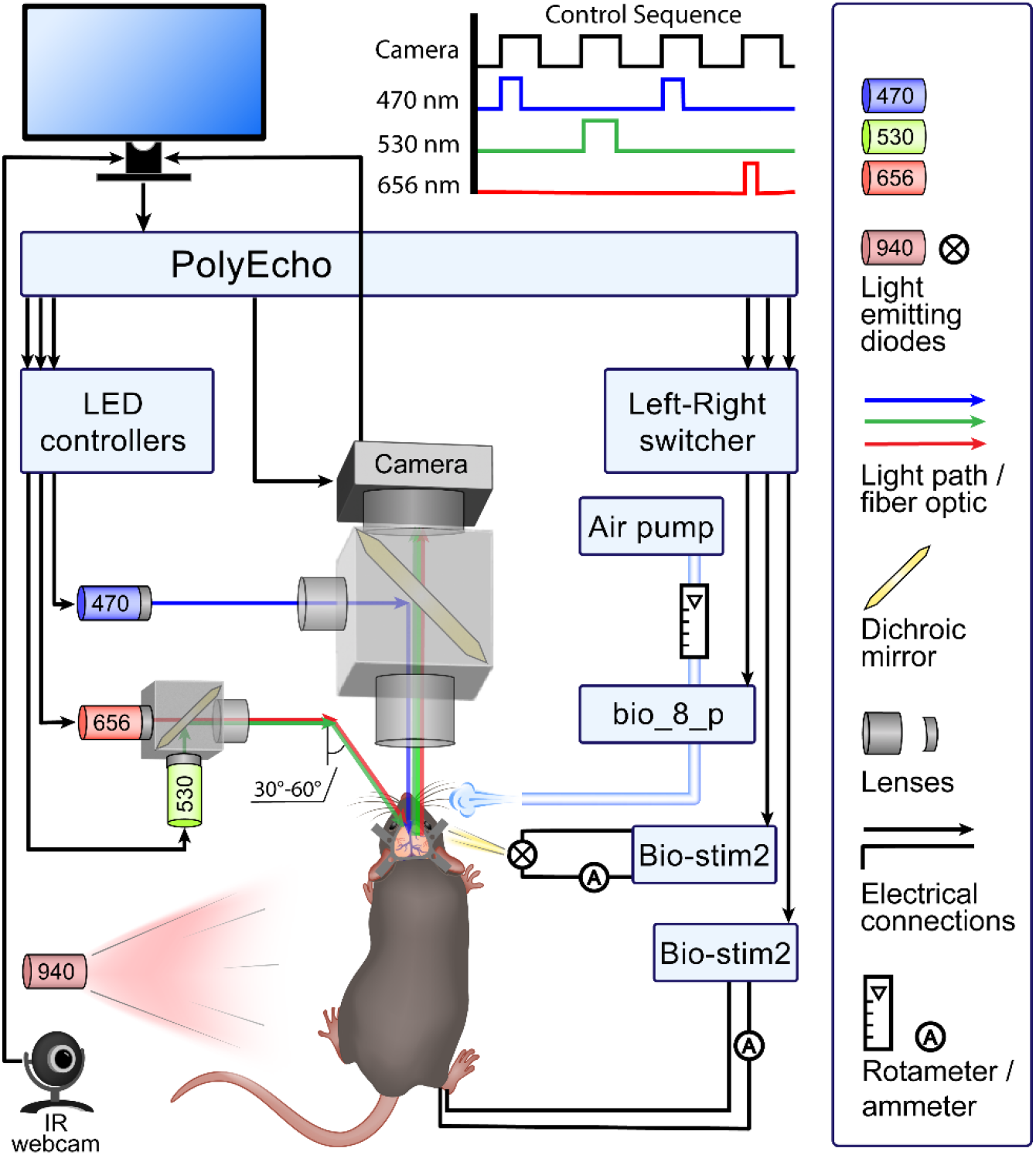
Schematic diagram of our WFOI system, with the illumination setup on the left and sensory stimulation devices on the right. An IR webcam with a 940 nm LED records mouse behaviour.

#### 3.1. The minimum configuration

The minimum configuration for a WFOI system includes:

- one LED for haemoglobin imaging;
- an LED controller to regulate current;
- a lens;
- a camera for image acquisition;
- an LED stand;
- a camera stand;
- two tables;
- a computer;
- software for controlling the camera and the LED controller;
- a multi-channel controller to synchronize the LED controller, the camera, and the computer;
- a head-fixation frame;
- a heating pad for the mouse.

This setup is enough for measuring Δ[HbT] or Δ[HHb] (depending on the chosen LED wavelength; see Table 2) in an anaesthetised animal.

**Table 2.** Decision-making for LED wavelength choice. The preferred wavelengths are highlighted with an underscore.

| Measured molecules | LED’s central wavelength | Reason |
| --- | --- | --- |
| HbT | 505 or <u>530</u> or 540 or 560 or <u>590</u> | Isosbestic points for HbO and HHb in the green-red spectrum are at 500, 530, 545, 570, and 584 nm. Use a narrow bandpass filter |
| HHb | <u>625</u>-680 | HHb has a 10× higher molar extinction coefficient than HbO in the red spectrum. Between 625 nm and 680 nm, HHb’s extinction decreases significantly |
| HbT, HbO, HHb | 590 and 625 nm | Longer wavelengths penetrate deeper into tissue, so yellow and red LEDs will sample similar depths (Ma et al., 2016). Theoretically, any two wavelengths are suitable. For optimal accuracy, one should be isosbestic, and the other should be in the red region |
| HbT, HbO, HHb, GCaMP | 530 and 625 nm for Hb<br>470 nm for GCaMP | 530 nm is near the emission maximum of GCaMP. Hb imaging is necessary for the correction of fluorescence signals.<br>430 nm may be used as a Ca-independent signal of GCaMP (not necessary) |
| HbT, HbO, HHb, FAD | 530 and 625 nm for Hb<br>470 nm for FAD | See above |
| HbT, GCaMP or FAD | 530 for HbT<br>470 nm for GCaMP / FAD | One wavelength near the emission maximum is enough for correction using the regression method (Valley et al., 2020) |
| HbT, HbO, HHb, ClopHensor | 530 and 625 nm for Hb<br>460, 505 and 560 nm for ClopHensor | See above |
| HbT, HbO, HHb, when intravenous dye is used (e.g., Rose Bengal) | E.g. 590 and 625 nm if Rose Bengal | Wavelengths outside dye absorption spectra |

Optional components that can enhance the system include: an uninterruptible power supply (UPS), an actively vibration-isolated table, a bandpass filter, collimating and focusing lenses, a light diffuser, and an external hard drive.

In our setup, the core system consists of a collimated LED 530 nm for HbT (BLS-LCS-0530-03-22, Mightex, CA, USA); a 530 nm bandpass filter (ZET532/10x, Chroma Technology Corp., VT, USA); a BioLED light source control module (BLS-1000-2, Mightex, CA, USA); lenses (OBJ-MAO-020-LWD, MAO-1-25-G-1C, Mightex, CA, USA); a high-sensitivity camera (CXE-B013-U, Mightex, CA, USA); a rectangular U-shaped LED mounting stand (Thorlabs, NJ, USA); an XYZ-motorised camera system (Zaber, Canada; Mightex, CA, USA), a U-shaped stand for motorized system (Thorlabs, NJ, USA; Micro Control Instruments Ltd., England); a custom-made anti-vibration table 75 × 270 cm and a secondary table for a mouse with a keyboard; a Back-UPS (ES 700VA, APS, USA); PolyScan4 and BuffCCD Cam software (Mightex, CA, USA); a Legion T5 26IOB6 computer (Lenovo Group Limited, China), and a GeForce 3060 Ti GPU (NVIDIA, USA); a 6 TB WDBWLG0040HBK-XB external drive (Western Digital, Thailand); a 12-channel PolyEcho Intelligent I/O Control Module BLS-IO12-U (Mightex, CA, USA); a Mobile HomeCage® (Neurotar, Finland), and a ThermoStar Homeothermic Monitoring System (RWD, China). The detailed description of mounting stands and computer characteristics are provided in Supplementary table S1.

*Note: Based on our experience, the camera, BuffCCD Cam software, the anti-vibration table, and the UPS included in the system are not optimal and may benefit from replacement. The equipment does not necessarily have to be expensive (see Discussion section), but a high-sensitivity camera is crucial for achieving a good signal-to-noise ratio*.

##### 3.1.1. Room, electrical supply and stands

1. Choose a dedicated recording room. Light, noise, and vibrations can interfere with imaging, therefore, cover windows with roller shutters. *Note: Temperature control in the room is important to prevent LED overheating*.
2. Set up a table for the LED, the camera and the head-fixation frame. An actively vibration-isolated table is recommended. Set up another table for the devices, that can cause vibrations (such as the computer). *Note: Ensure that monitor light does not reach the experimental setup. Place the monitor behind the mouse to avoid unintended visual stimulation*.
3. All critical equipment should be backed up with a UPS of appropriate VA capacity (an online UPS is optimal). Connect the UPS (or multiple units if required) to outlets on separate electrical circuits.
4. Install the head fixation stand (e.g., a stereotaxic frame). Assemble the LED mounting stand and the camera stand. *Note: At least one of the following should be movable to allow proper focusing: the camera stand or the head frame. Tighten all screws securely to minimise vibrations. For further vibration control, consider fixing the stands to the tabletop. If the frame shifts by more than a pixel, post-recording alignment will be required (see 5*.*1)*.

##### 3.1.2. Optical system

1. Assemble the LED for light scattering on its stand to provide lighting at an angle of 30–60°. We recommend LED models with passive cooling and a central wavelength of 530 or 590 nm for HbT and 625 nm for HHb. The use of one LED will not allow precise measurements as HbO and HHb spectra overlap (Fig. 5). To measure the Δ[HbT] and Δ[HHb] precisely see 3.3.3.1 and 5.2.2. *Note: Lighting at a 90° angle for scattering creates glare from the skull surface, interfering with brain activity observations*.
2. Add a bandpass filter of ~20 nm bandwidth after the LED to improve measurement accuracy. The recommended LED output power is 200–500 mW when a narrow bandpass filter is used. *Note: Lower-power LEDs or optical filters with narrower bandwidths may require longer exposure times (10–20 ms) — ensure this is acceptable for your experiment*.
3. Mount collimating lenses to direct LED light. This reduces light loss. *Note: LED light may act as an unintended visual stimulus for the mouse*.
4. Connect the LED to the LED controller. The controllers must automatically trigger the LED and manage the electrical current (not voltage) supplied to the LED to regulate brightness. Always consider the maximum allowable current for the LED. *Note: Use LEDs and LED controllers from the same manufacturer*.
5. Set up a light diffuser between the LED and the mouse. You can make it from glass or PET plastic sanded with sandpaper. Another option is to use a circular polarizing filter mounted on the lens, but it reduces the overall image brightness by 75% or more. *Note: A diffuser helps eliminate glare. Surface irregularities of the skull become more noticeable, but this does not hinder the recorded brain activity*.
6. Select a camera. We recommend using a monochrome camera with external trigger control (TTL signal), a frame rate of 100 Hz at a resolution of at least 256 × 256 pixels, and low dark current. However, a maximum frame rate of 10 Hz is sufficient for HHb- or HbT-only measurement. (If the camera’s dark signal is not zero, increase the frame rate to capture dark frames; if sensitivity is low, increase the frame rate for subsequent temporal binning.) The captured images should have a bit depth of at least 16 bits. *Note: 8-bit images can also be used, but the processing quality will be lower. Resolution is not critical; for WFOI analysis binning down to 64 × 64 pixels can be used. A camera with global shutter is preferable*.
7. Attach the camera to the lens and set it up on the camera stand above the animal’s head. A 2× – 4× lens with a working distance of 10 cm is recommended.

**Figure 5.**
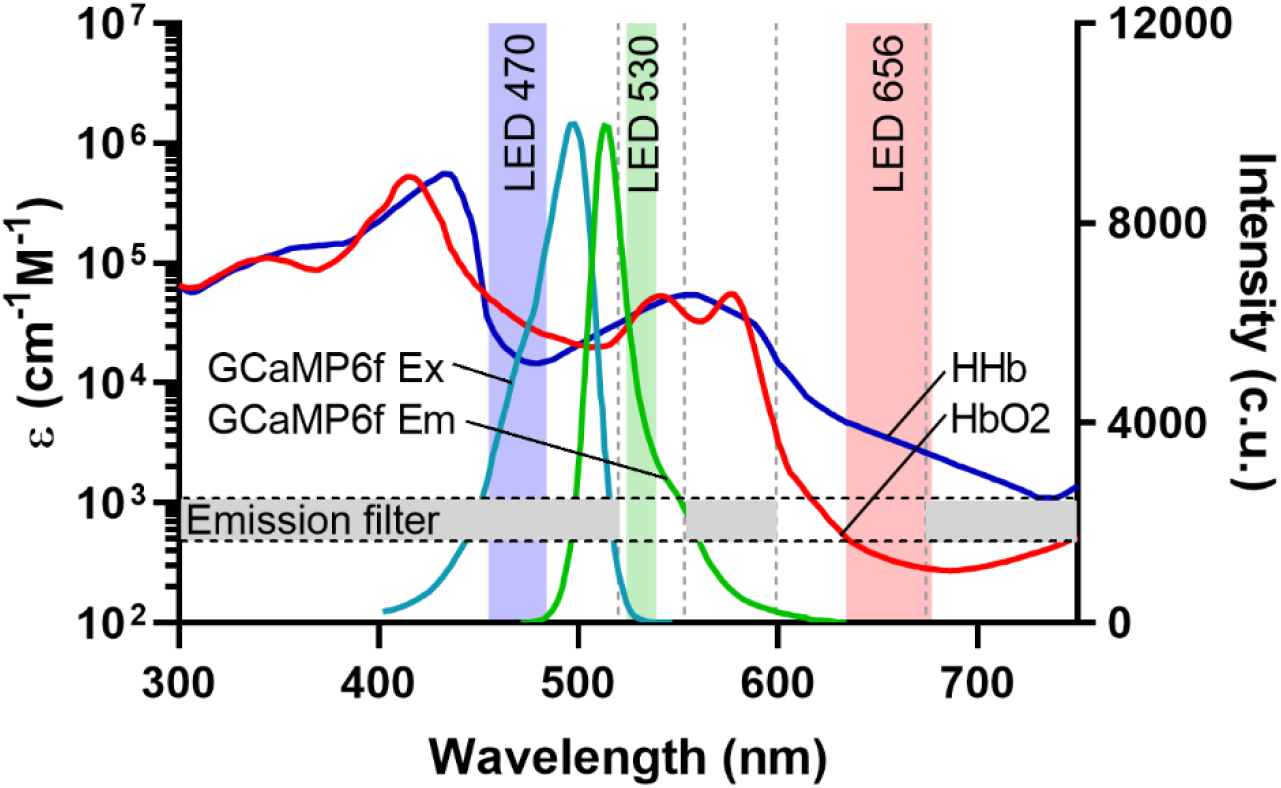
Optical spectra of our WFOI setup for the measurement of oxyhaemoglobin (HbO) and deoxyhaemoglobin (HHb) alongside Δ[Ca^2+^]_i_. The figure shows the molar extinction coefficients for HbO and HHb, and the excitation and emission spectra of GCaMP6f. LED spectra are shown after bandpass filers. The white sectors on the “Emission filter” stripe correspond to the transparency windows of the emission filters. The same configuration can be used for Δ[FAD] measurements in wild-type mice (not shown).

*Notes:*

- *A lens with a shorter working distance will capture more signal but will make mouse manipulation more difficult*.
- *Although optional, steps 2,3, and 5 markedly enhance imaging quality*.

##### 3.1.3. Control system

1. Connect the LED controllers to an analog output of a multi-channel controller. Connect the camera TTL trigger to a digital output. The multi-channel controller will transmit signals from the computer. The number of required channels depends on the number of wavelengths used — one per wavelength, one for the camera, and one per external device (e.g., a stimulator).
2. Connect all devices to a computer or laptop. The computer should meet the following minimum system requirements: 16 GB of RAM, 512 GB of storage (preferably SSD), an Intel Core i5 or equivalent processor, and Windows 10/11 (64-bit), USB 3.0 support (at least four ports). *Note: For optimal performance, we recommend a computer with 64 GB of RAM, an Intel Core i7 (or equivalent) processor, and a dedicated graphics card (GPU). Pay attention to the number of USB ports: in a full setup (see the following sections), eight ports will be occupied*.
3. Install all necessary software for controlling the LEDs and the camera. This can be commercial software provided with the multi-channel controller and the camera or a third-party software (e.g. Micro-Manager 1.4 for the camera (D. Edelstein et al., 2014)). The software should be capable of starting and stopping recordings, controlling the activation, intensity, and duration of LED illumination, adjusting camera exposure time, synchronizing frame exposure with LED activation, and configuring image-saving parameters (binning, gain, file format, automatic file naming, save path). Additionally, it should allow pre-recording adjustments, including a preview mode for camera focusing and LED duration selection.
4. For raw data storage, select an external hard drive (6–8 TB).

#### 3.2. LED illumination modules and lightguides

The combination of LEDs in LED illumination modules is advisable for multi-wavelength illumination (see below). Liquid light guides eliminate the need for LED placement in close proximity to the mouse’s head; they also improve the illumination angle and radius. Grouping the LEDs simplifies data processing later due to a uniform illumination pattern.

In our system, we combined 530 nm and 656 nm LEDs with 1-to-7 fibre optic bundle in one module, and 460, 505, and 560 nm LEDs with a single-fibre light guide in another module. Particular components depend on LED wavelength; please see 3.1, 3.2, 3.4, 3.5 for details.

1. Select appropriate beam splitters to combine LEDs for haemodynamic measurements into one assembly and those for fluorescence into another. Arrange the LEDs in the assembly in order of increasing wavelength (the shortest wavelength closer to the output of the optical fibre). Ensure that each LED achieves >95% transmission or reflection. *Note: Alternatively, use one LED assembly for both haemodynamics and fluorescence (for bright sensors)*.
2. Assemble the LED illumination modules: attach collimating lenses (see 3.1.2.3) to the LEDs, place bandpass filters (see 3.1.2.2 and 3.4.2) into the designated recesses on the beamsplitter, and affix each LED to the beamsplitter. Connect a lightguide adapter with focusing lens to the beamsplitter’s output. *Note: Follow the manufacturer’s recommended light direction for the dichroic mirrors. If LED power is suboptimal, use neutral density (ND) filters along with bandpass filters*.
3. Connect a light guide to a lightguide adapter. The selected light guide must be long enough to allow camera movement. For fluorescence, we recommend a 1-metre-long, 3-mm core diameter liquid light guide. For haemodynamics we recommend 1.2 m-long branched light guide (1–7) with 0.6-mm core diameter. *Note: Handle the light guide carefully to avoid end scratching the ends and do not exceed the minimum bend radius. With high-power LEDs, it can be replaced by inexpensive optical fibre*.
4. Adjust the light guide at the proper angle to the plane of the headplate: 30-60° for haemodynamics, 90° for fluorescence (see 3.1.2.1, 3.2.1 note, and 3.4.2). It is optimal to adjust the fibre optic bundle for haemodynamics around the lens (see 3D model for the mount in Supplementary file S3) to prevent shadow formation.

#### 3.3. HbO, HHb and HbT measurements

HbO and HHb have different absorption spectra (Fig. 5). Two LEDs of wavelengths with distinct molar extinction coefficients are enough to calculate both Δ[HbO] and Δ[HHb] (Ma et al., 2016).

Our HbO/HHb measuring system includes the core system (see 3.1) and the following components: an LED at 656 nm (LCS-0656-07-22, Mightex, CA, USA), a bandpass filter (AT655/30m, Chroma Technology Corp., VT, USA), a beam splitter (LCS-BC25-0605, Mightex, CA, USA), a collimating lens with a lightguide adapter (LGC-019-022-05-V, Mightex, CA, USA) a 1-to-7 fan-out fibre optic bundle (BF76HS01, Thorlabs, NJ, USA), and a custom-made mount for fibre optic bundle (see Supplementary file S3).

1. Select the second LED for haemodynamic imaging based on the haemoglobin absorption spectra (Fig. 5). For intrinsic signal imaging only, we recommend 590 nm and 630 nm. If simultaneous GFP-based protein imaging is used, we recommend 530 nm and 630 nm (Table 2, see also 3.1.2.1).
2. If available, assemble the LED illumination module with a fibre optic bundle and a mount for shadow-free illumination (see 3.2).
3. Connect the second LED to the LED controller. If you have a 2-channel controller with the proper maximum current, use it for both LEDs.
4. Consider the frame rate (see 3.1.2.6), the number of channels in the multi-channel controller (see 3.1.3.1), and the software requirements (see 3.1.3.3).

#### 3.4. FAD and Ca^2+^ measurements

Due to the strong overlap between the excitation and emission spectra of FAD and GCaMP, fluorescent measurements of both Δ[FAD] and Δ[Ca^2+^]_i_ can be performed using the same optical setup. Δ[FAD] can be measured in wild-type mice (e.g., C57BL/6J, IGB RAS, Russia), whereas Δ[Ca^2+^]_i_ can be measured in transgenic animals, e.g., strain GP5.17 (JAX, stock #025393). FAD fluorescence is weak, and FAD concentration changes are of low amplitude, and therefore make a minor (4-11%) contribution to Ca^2+^ measurements with GCaMP (Kozberg et al., 2016).

Fluorescent measurements require simultaneous measurements of Δ[Hb], as dynamically changing Hb absorbs both excitation and emission light (Ma et al., 2016; Valley et al., 2020).

In our setup, fluorescent measurements of FAD and GCaMP are performed using an LED at 470 nm (BLS-LCS-0470-14-22-J, Mightex, CA, USA), a bandpass filter (ET470/24m, Chroma Technology Corp., VT, USA), a BioLED Light Source control module (BLS-3000-2, Mightex, CA, USA), a liquid light guide (#77566 3-mm diameter, 1-m length, Thorlabs, NJ, USA), collimating lenses (LGC-019-022-05-V – 2 pcs, EPI-MAO-LG, Mightex, CA, USA), and a fluorescent cube (CUBE-MAO-CARR, Mightex, CA, USA) with filters (59009bs, 59026m, Chroma Technology Corp., VT, USA).

1. Select an LED of the appropriate excitation wavelength (470 nm for GCaMP, 460 or 470 nm for FAD). Use (*FPbase*, 2025) to determine the optimal wavelength (Table 2). The required LED power depends strongly on the fluorophore brightness, the optical filters’ bandwidth, the camera sensitivity, the exposure time, and the possibilities for temporal binning. In our setup, an output power of 860 mW is suitable for both FAD and GCaMP measurements using spatial binning.
2. Assemble a fluorescence filter cube. Mount a bandpass optical filter facing toward the LED, a dichroic mirror at a 45° angle, and an emission filter facing toward the camera. These components must prevent excitation light from entering the camera while efficiently transmitting light in the emission range. For examples, see (*Chroma Technology*, 2025). *Note: The cube dimensions determine the filter diameter and, therefore, overall system cost*.
3. Place a collimating lens between the LED (or the light guide, if used; see 2.2) and the filter cube to transform the divergent light into a parallel beam. *Note: An inappropriate lens may create an interference pattern. Processing of such frames is complicated, but possible*.
4. Shield the LED light sources. Seal any gaps through which light could escape to prevent stray light from influencing the recordings.
5. Consider the frame rate (see 3.1.2.6), the number of channels in the multi-channel controller (see 3.1.3.1), and the software requirements (see 3.1.3.3).

#### 3.5. Cl- and pH measurements

In transgenic mice expressing ClopHensor (Arosio et al., 2010; Diuba et al., 2020b; Ponomareva et al., 2021) ΔpH_i_ with Δ[Cl^−^]_i_ can be measured via fluorescence. The setup includes an LED illumination module with three LEDs for excitation (BLS-LCS-0455-03-22, BLS-LCS-0505-12-22, BLS-LCS-0560-03-22, Mightex, CA, USA), a BioLED Light Source control module (BLS-1000-2, Mightex, CA, USA), bandpass filters (#65-142, Edmund Optics, USA; ET505/20x, ZET561/10x, Chroma Technology Corp., VT, USA), and beam splitters (LCS-BC25-0480, LCS-BC25-0515, Mightex, CA, USA). The other parts remain unchanged, as we select filters for the fluorescence cube as universal for all three fluorophores (FAD, GCaMP6f, and ClopHensor). A 460 nm LED with a narrow bandpass filter is crucial to reach an isosbestic point for pH and therefore to measure Δ[Cl^−^]_i_.

#### 3.6. Recordings in an awake animal

Anaesthesia significantly affects brain activity, although both the recording procedure and data processing in an awake animal are complicated.

Our setup consists of the Mobile HomeCage® (Neurotar, Finland), a custom-made cage with transparent walls, two C-110 63110 webcams with removed IR filters (Defender, China), and two Helios IR-Plate-2-940 illuminators (Microlight, China).

*Note: Based on our experience, the C-110 63110 webcam is not optimal and may benefit from replacement*.

1. Select an appropriate head fixation system, ensuring that it is safe for the mouse. The fixation system should be comfortable, adjustable in height, quick to secure, and should not interfere with exploratory behaviour.
2. Set up a movable platform under the head fixation system. This could be an air-supported cage, a treadmill, a rotating disk, or a spherical ball. *Note: An awake animal requires a system that permits running (preferably, in all directions) and grooming without risking head injury. An air-supported cage with opaque walls can create an illusion of free movement; however, we recommend using transparent walls to enable behavioural video recording*.
3. Mount the necessary components for behavioural video recording, including at least one infrared (IR) illuminator and one or more cameras. *Note: Behaviour significantly affects brain activity (see Fig. 10); therefore, it is essential to segment the recording into active and resting periods. Any spectrum outside the range captured by the brain imaging camera may be used; however, IR light is particularly suitable, as it is invisible to mice and does not introduce an additional sensory stimulus*.
4. Set up additional components, such as an airflow control unit and a pump.

#### 3.7. Sensory stimulation

Sensory stimulation in an awake animal is compatible with WFOI. Behaviour can interfere with the brain’s response to stimulus, whereas anaesthesia dramatically reduces brain activation. Sensory stimulation facilitates functional cortical mapping both under physiological conditions (e.g., throughout development) and in disease models (e.g., stroke), and allows for evaluation of the impact of external or experimental factors on fundamental brain functions.

Our stimulation system for the eyes and hind limbs includes (Fig. 6): an isolated constant current electrical stimulator (Bio-stim2, Biotechnologies, Russia); a microammeter; a 3mm white LED; a custom-made left-right switcher; and custom-made hind limb electrodes with a tail clip.

**Figure 6.**
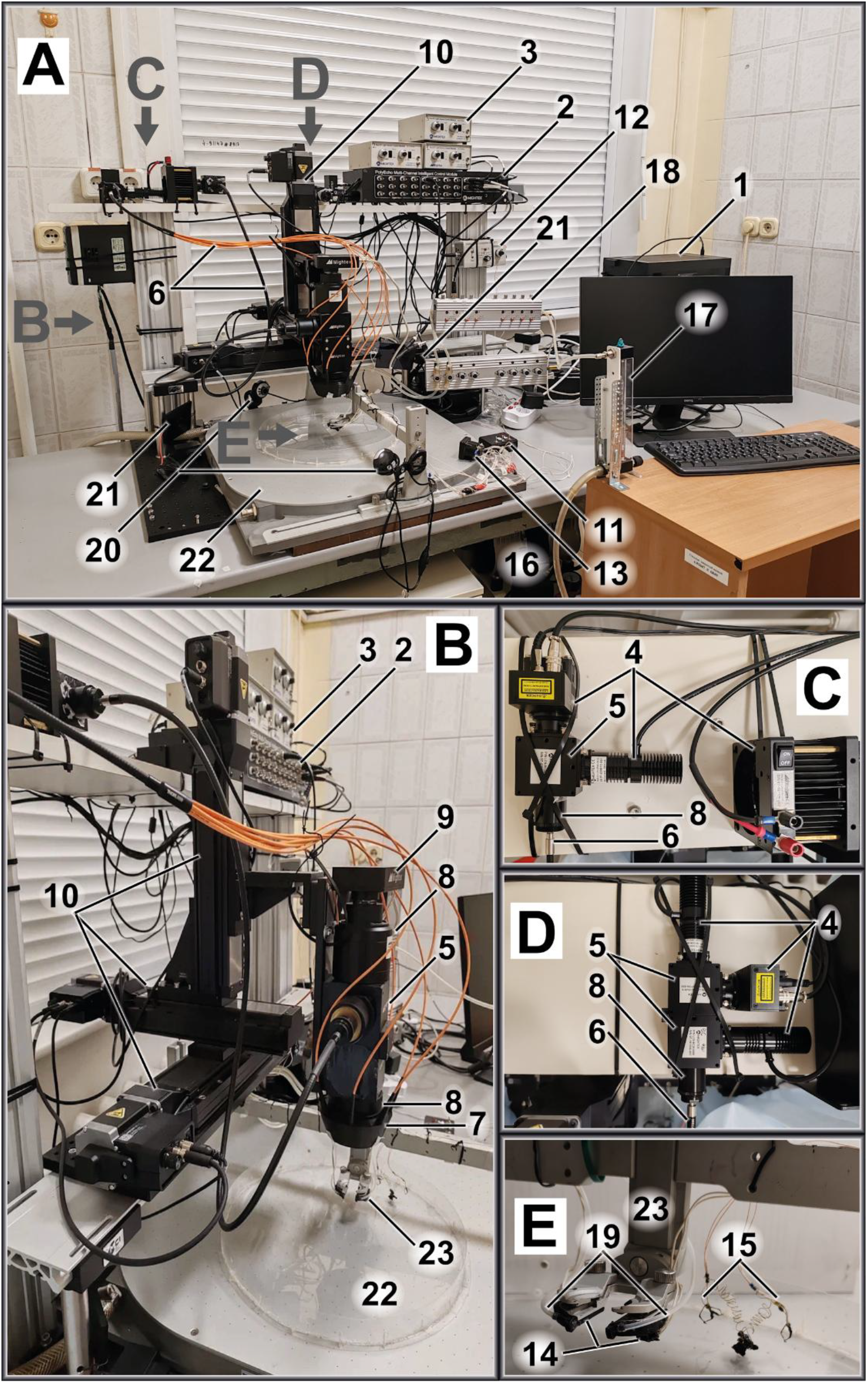
The whole WFOI setup. Panels B, C, D, and E show more detailed views of the setup in the planes indicated by the arrows in A. <u>Control system</u>: 1 - Computer; 2 - PolyEcho; 3 - LED controllers. <u>Brain lighting and recording system</u>: 4 - LEDs with bandpass filters; 5 - Cube with filters and dichroics; 6 - Fibre optics (1-to-1 and 1-to-7); 7 - 1-to-7 fibre optic mount; 8 – Lenses; 9 - CCD camera; 10 - XYZ camera positioning. <u>Eye and limb stimulation:</u> 11 - Left-right switcher; 12 - Electrical stimulator; 13 – Ammeter; 14 - White LED; 15 - Hind limb electrodes with tail clip. <u>Vibrissae stimulation:</u> 16 - Air pump; 17 - Rotameter; 18 – bio_8_p (air tubes pinching device); 19 - Air tube. <u>Mouse behaviour:</u> 20 - Webcam with extracted IR-filter; 21 - IR LEDs. <u>Mobile HomeCage:</u> 22 - Mobile HomeCage® perforating air-dispensing platform (air pump not shown) with a custom-made cage with transparent walls; 23 - Mobile HomeCage® head-fixation apparatus.

Our stimulation system for vibrissae includes: an air pump (607CD22, Thomas Industries Inc., USA); silicone air tubes with diameters of 10, 7, and 2 mm; a rotameter (RWD, China); and a device for pinching air tubes (bio_8_p, Biotechnologies, Russia).

White LEDs and air tubes are mounted on a 3D-printed attached to the head fixation frame. We found it optimal for the LED light to be directed downward. Hind limb electrodes are clip-like and are secured to a tail clip with two soft springs. During recording, the electrodes are attached to ring-shaped piercings (two rings per limb, hooked onto the Achilles tendon). The piercings were fabricated from 27G needles; the electrodes were made from segments of a second guitar string; and the tail clip was constructed from a hair claw clip.

### 4. Head-fixation training and imaging procedure

This section describes animal habituation to the head-fixation setup, parameter selection, and the imaging procedure. Since setup configurations and experimental objectives may vary across laboratories, we aim to outline general principles rather than focus on the specifics of our setup.

#### 4.1. Mouse habituation to head fixation

The habituation protocol acclimatises the mouse to head fixation in several stages. On day one, the mouse is familiarised with the room for 1h. The mouse is then gently restrained 3*–*5 times by the head in the handler’s hands, then placed on a movable platform near head-fixation apparatus for 1 min. On day two, the same steps are repeated, with addition of mouse restraining for short-term in the setup. On day three, the steps from day two are repeated, with the addition of a longer head fixation (approximately one minute) in the apparatus, ending with the lights off and the LEDs on. Each manipulation is repeated at least three times, with rest periods in between. The aim is for the mouse to remain calm during fixation, without associating the setup with negative experiences (for the detailed SOP, see Supplementary file S2; for troubleshooting guide, see Supplementary file S4.).

*Notes: Mice should be well acclimatised to handling before starting. Ensure the room’s noise level matches experimental conditions*.

#### 4.2. Selection of imaging parameters

1. Select parameters according to your experimental goals. Choose a recording duration of 3– 5 min for resting-state imaging or longer for stimulation protocols (at least 20 stimuli should be presented at 20-second intervals). Set the frame rate based on signal dynamics: ≥20 Hz for calcium imaging, and 10 Hz for slower signals (Hb, FAD, Cl^−^, and pH). Define the LED sequence appropriate to the indicators used (e.g., LED470 → LED530 → LED470 → LED656 for GCaMP, HbT, and HHb).
2. Configure the imaging system and triggering. Set up the software (e.g., PolyScan4) according to the manufacturer’s instructions, including LED models and controllers. Programme an external TTL trigger matched to the frame rate with pulse duration shorter than the frame interval (e.g., 1 ms), and set the camera exposure time directly in the camera settings so that the pulse duration does not affect it. The number of pulses = frame rate (Hz) × recording duration (s).
3. Set camera and exposure parameters. Use a fixed camera exposure time (e.g., 5 ms) for all recordings to avoid aliasing. Configure the camera to acquire 16- or 32-bit TIFF or RAW images at a resolution of 256 × 256 pixels (with binning), with gain set to 0. Enable timestamp-based file naming and continuous preview mode.
4. Calibrate exposure and verify system performance. Place a 1 × 2 cm piece of white paper with blue, green, and red markings in the field of view. Focus the camera, then gradually increase the camera exposure for each LED until glare appears; set the final LED exposure to 80 % of the glare threshold. If exposure values differ significantly, reduce intensity of the brightest LED (not below 50 % of maximum current, use neutral density filter if needed). Align LED pulse onset with camera triggers, use LED exposure, defined previously. Restore the fixed camera exposure time and enable trigger mode. Run a test acquisition to confirm:
  a. Even illumination and wavelength-specific disappearance of coloured marks
  b. No image shift (≤1 pixel)
  c. Absence of interference from other devices (test during activation)
  d. Stable ROI intensity traces
5. Save your configuration. Archive all camera and system settings to ensure consistent application in future recordings.

#### 4.3. Imaging procedure

1. Prepare the equipment and protocol. Power on all devices and the control computer. Load the imaging protocol and camera settings. Set the save path and enable automatic file naming.
2. Acclimate the animal. Transfer the mouse to the imaging room and allow 30 minutes for habituation.
3. Position and prepare the animal. If required, induce and maintain anaesthesia, turn on the heating pad, and apply ophthalmic gel. Fix the mouse securely in the setup, clean the skull, and protect the eyes with a visor (see 1.1.4). *Note: Hand-tightening of the holding screws for mouse fixation may be insufficient*.
4. Set up the imaging system. Focus the camera, turn on one LED in continuous mode, and switch off ambient lighting.
5. Adjust exposure and LED timing. Optimise exposure for each LED to 70–80 % of the glare threshold (see 4.2.4). Restore the selected exposure and adjust the LED pulse durations. Switch to trigger mode.
6. Start recording and minimise disturbance. Begin imaging. Remain quiet and still to avoid external stimulation. Limit total head-fixation time to 1–2 hours.
7. Finish the session and store data. Return the mouse to its home cage and transfer the raw data to external storage.

### 5. Image analysis

Wide-field optical imaging produces sequences of thousands of images. Extracting meaningful signals requires image alignment, noise reduction, and the conversion of pixel intensity into relative changes in haemoglobin concentration or sensor fluorescence. Fluorescent signals must also be corrected for haemoglobin absorption. Additional challenges include automation, faster analysis, and efficient data storage. While various research groups have developed WFOI analysis pipelines in MATLAB and Python, existing open-source scripts lack universality due to variations in hardware, imaging protocols, and analysis methods (Brier & Culver, 2023; Haupt et al., 2017; Xiao et al., 2021). To address this, we developed a Python-based software package tailored for our WFOI setup, described below and available on GitHub (https://github.com/natalia1896/WIFIOPIA). The key stages of the programme’s operation are listed below.

#### 5.1. Image annotation and brain masking

Due to manual headplate implantation, the accessible cortical area and the implantation angle may vary slightly across animals. Additional variability may arise from small shifts in objective positioning between imaging sessions. To enable cross-subject comparisons, images are spatially aligned using anatomical landmarks such as bregma and lambda. These landmarks are manually annotated at the start of analysis, and the corresponding rotation and translation parameters are applied to standardise image orientation across sessions. A brain mask is also required to define cortical regions for analysis. This mask can be generated manually or via automated thresholding or neural network-based segmentation. In our workflow, we use manual annotation via a custom graphical interface, though externally generated binary masks are also supported.

#### 5.2. Image transformation and physiological signal conversion

##### 5.2.1. Image transformation

Based on the annotated landmarks, all images are spatially aligned and sorted into 3D arrays (t, w, h), where each array corresponds to a specific excitation wavelength defined by the illumination protocol. Signal changes are then computed as relative intensity values (I/I_t=0_), where I_t=0_ represents either the mean signal over the full acquisition or over a selected baseline period.

##### 5.2.2. Scattering signal conversion to haemoglobin concentrations

For dual-wavelength reflectance recordings, relative changes in haemoglobin concentrations (Δ[HbO], Δ[HHb], Δ[HbT]) are calculated using the modified Beer–Lambert law, as described in (Ma et al., 2016):

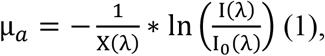

where μ_*a*_ is the absorption coefficient, X(λ) is the wavelength-dependent pathlength. In our experiments, we use wavelength-dependent pathlengths from (Ma et al., 2016). Another step is to solve the following system of equations:

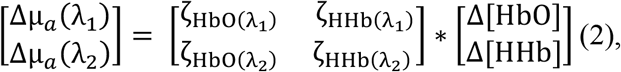

where ζ_Hb(*λ*)_ is the molar absorption coefficient of specific form of haemoglobin at wavelength λ, Δ[HbO], Δ[HHb] are the changes (relative to baseline I_t=0_) in the molar concentrations of HbO and HHb, respectively, Δμ_*a*_(*λ*) – the changes (relative to baseline I_t=0_) in absorption coefficients.

We use tabulated molar extinction coefficient for haemoglobin in water, as provided by (*Oregon Medical Laser Center*, 2025). To calculate ζ_Hb(*λ*)_ from the tabulated data, the following equation was used:

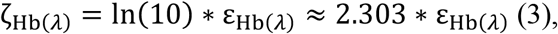

where ζ_Hb(*λ*)_ is the molar absorption coefficient (in natural logarithm form), and ε_HbO(*λ*)_ is the decadic molar absorption coefficient. Both used in the Beer-Lambert law, either in its exponential form 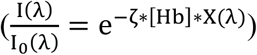, or in its absorbance form 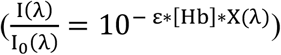.

The final solution to the equation is presented below:

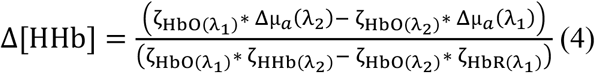

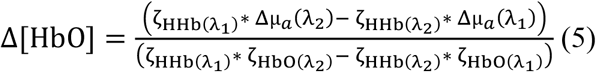

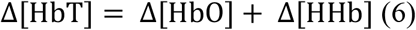

When only one reflectance wavelength is used, and it lies within the isosbestic range (e.g., 530 nm), changes in ΔI/I_t=0_ (%) can serve as a proxy for total haemoglobin fluctuations.

##### 5.2.3. Correction of fluorescence signal for haemoglobin absorption

At present, the search for optimal fluorescence correction methods remains a methodological challenge in the field. Here, we describe two such approaches.

Our code includes an implementation of haemoglobin correction based on the modified Beer– Lambert model (Ma et al., 2016):

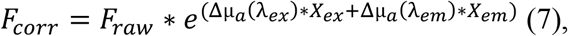

where F_raw_ is the raw F/F_0_, *X*_*ex*_ and *X*_*em*_ are photon pathlengths at excitation and emission wavelengths, respectively. However, this approach requires knowledge of the modified excitation and emission photon pathlengths (*X*_*ex*_, *X*_*em*_), a parameter that is often experiment-specific and difficult to measure directly (Sunil et al., 2023).

We also implement a data-driven correction using pixel-wise least squares regression (Valley et al., 2020). For each pixel (y, x), a linear model is fit to the observed fluorescence F_raw_(t) using the corresponding haemodynamic trace Hb(t) (e.g., Δ[HbT] or ΔI/I_t=0_) as a predictor:

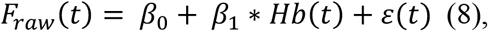

where *β*_*0*_ and *β*_*1*_ are the coefficients, estimated by the linear model, and ε(t) represents the residuals.

The regression is solved using `np.linalg.lstsq`, minimising the squared residuals. The corrected fluorescence signal is then represented by the residuals after removing the haemodynamic component:

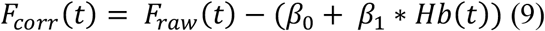

In our experiments, pixel-wise least squares regression demonstrated the best results.

The final output includes percentage changes in fluorescence (ΔF/F_0_) and haemoglobin concentrations in µM (Δ[HbO], Δ[HHb], Δ[HbT]), or relative reflectance changes (ΔI/I_t=0_, %) for single-wavelength recordings. Optionally, the global signal (mean intensity across the brain mask) can be subtracted to reduce fluctuations caused by illumination variability. Additional post-processing steps, such as Gaussian smoothing or frequency filtering, can be applied to minimise noise and improve signal quality.

## Results

The described protocol for creating a wide cranial window provides an opportunity to accurately measure brain haemodynamics without the need for a craniotomy (Fig. 7). During the thinning process, diploic vessels are removed; therefore, haemodynamics unrelated to neuronal activation does not obscure the underlying activity. Removal of the cancellous bone also improves image quality, as it contributes substantially to light scattering.

**Figure 7.**
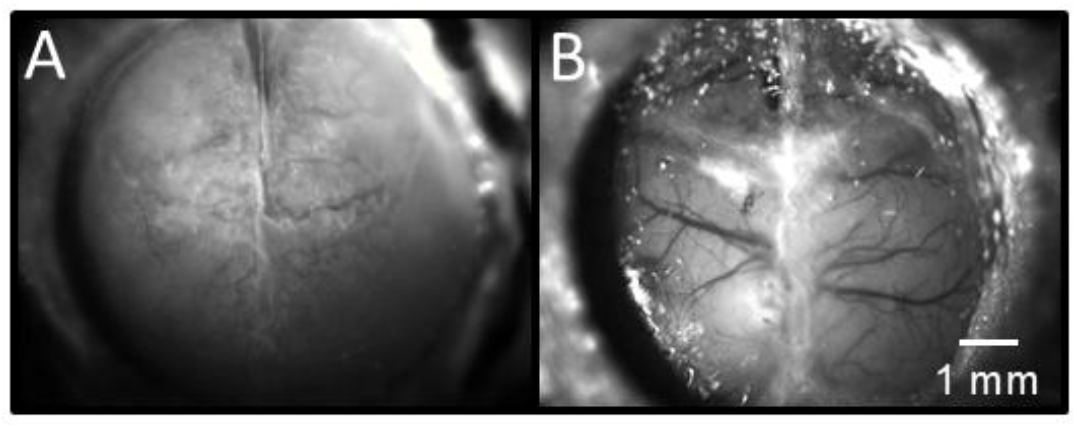
Representative images of the unthinned (A) and thinned (B) scull at 530 nm in light-scattering (reflectance) mode. During the thinning process, cancellous bone containing diploic vessels is removed.

The skull thinning technique provides optical access for recording cortical activity in the same animal for at least three months (Fig. 8). The durability of the cranial implant ensures stable recordings in awake animals during chronic experiments, withstanding multiple fixation and de-clamping procedures.

**Figure 8.**
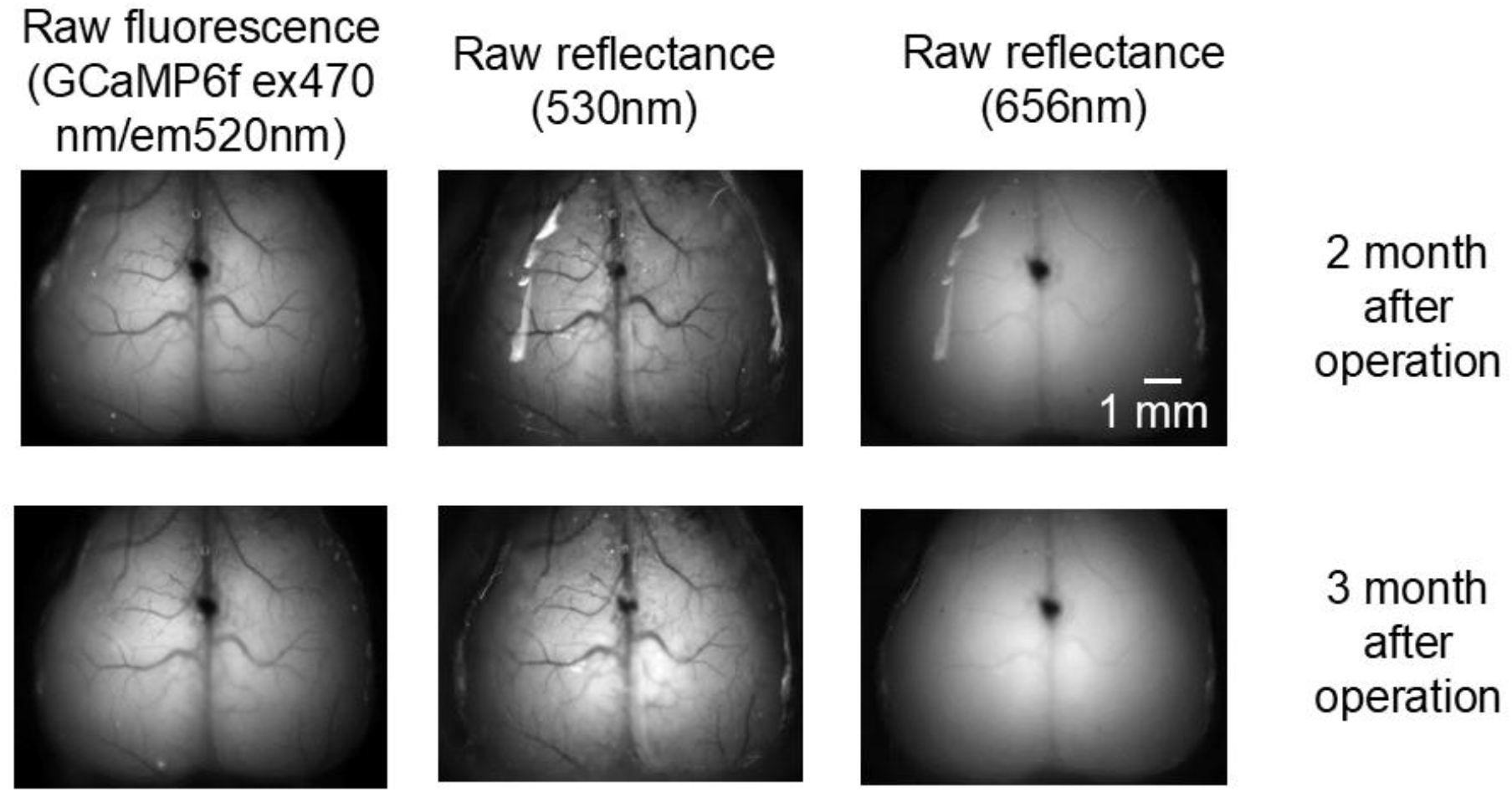
Representative raw images of the cranial window. The proposed protocol for creating a wide cranial window enables stable optical access to the mouse cerebral cortex for over three months. Shown are representative raw fluorescence images from the GCaMP6f sensor and light-scatter images acquired under 530 nm and 656 nm illumination, taken two and three months after surgery from the same animal.

The wide cranial window enables the recording of activity across extensive cortical regions in mice, including the motor, somatosensory, parietal, and retrosplenial cortices. To validate the applied brain activity recording and image analysis approaches, we assessed cortical responses to sensory stimulation of different modalities (Fig. 9).

**Figure 9.**
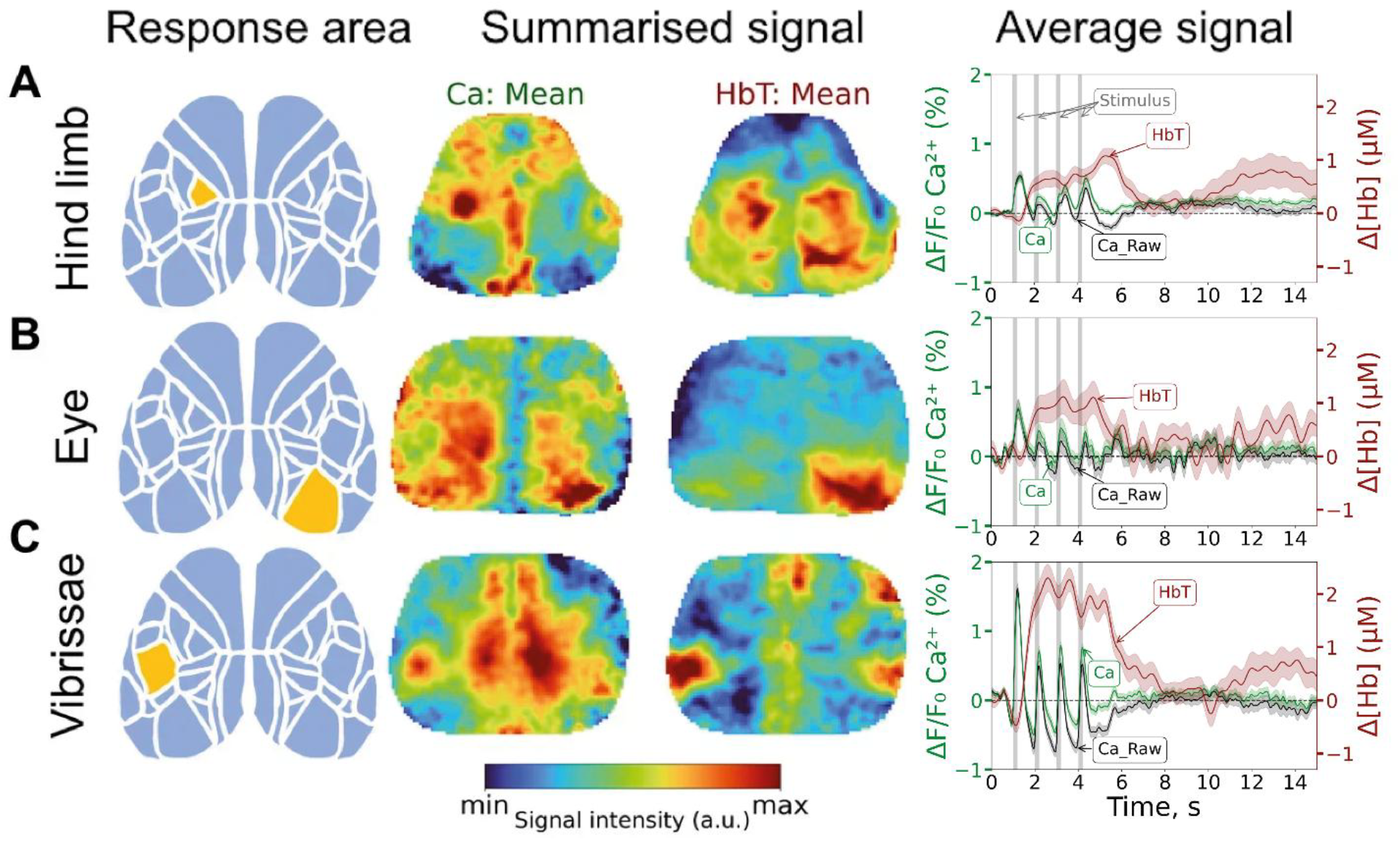
WFOI enables the recording of cortical responses evoked by sensory stimuli of different modalities. Responses to stimulation of the right hind limb (**A**), the left eye (**B**) and the right vibrissae (**C**). From left to right: (1) the location of the corresponding response area according to the anatomical atlas (Q. Wang et al., 2020); a summarised calcium (2) and haemodynamic (3) signal during the presentation of the stimulus, normalised from maximum to minimum over 21 repetitions; (4) the average calcium signal from the response area before and after haemodynamic correction (black and green lines, respectively), along with the haemodynamic response (dark red line).

We successfully recorded localised cortical responses to paw, whisker, and eye stimulation. The spatial distribution of activation areas closely matched the anatomically defined sensory regions for each modality, as referenced in standard brain atlases. Stimulation evoked an increase in the calcium signal, followed by a localised haemodynamic response indicative of blood influx.

A stimulation protocol comprising 21 trains of four short, repetitive stimuli (200 ms duration, 1 Hz frequency) was employed. This design enabled precise localisation of the response areas.

Importantly, stimulus-evoked responses in awake animals are affected by behavioural state (Fig. 10). The integral measure of calcium response – area under the curve (AUC) – was significantly lower when animals remained behaviourally calm (0.9 ± 1.11 [mean ± SD], n = 146), compared to trials involving active locomotion (3.16 ± 2.83, n = 232; Welch’s t-test: t(326.4) = –10.90, p < 0.0001; Cohen’s d = –0.97). A more pronounced effect of movement was observed on the haemodynamic response (measured 0–10 s after stimulation start): AUC of Δ[HbT] was 5.11 ± 15.20 (n = 146) during calm responses and 48.91 ± 50.14 (n = 232) during locomotion (Welch’s t-test: t(293.4) = –12.43, p < 0.0001; Cohen’s d = –1.08). Additionally, behavioural activity increased the variability of both calcium and haemodynamic signals, as indicated by Levene’s test (calcium: F = 71.67, p < 0.0001; haemodynamics: F = 173.63, p < 0.0001).

**Figure 10.**
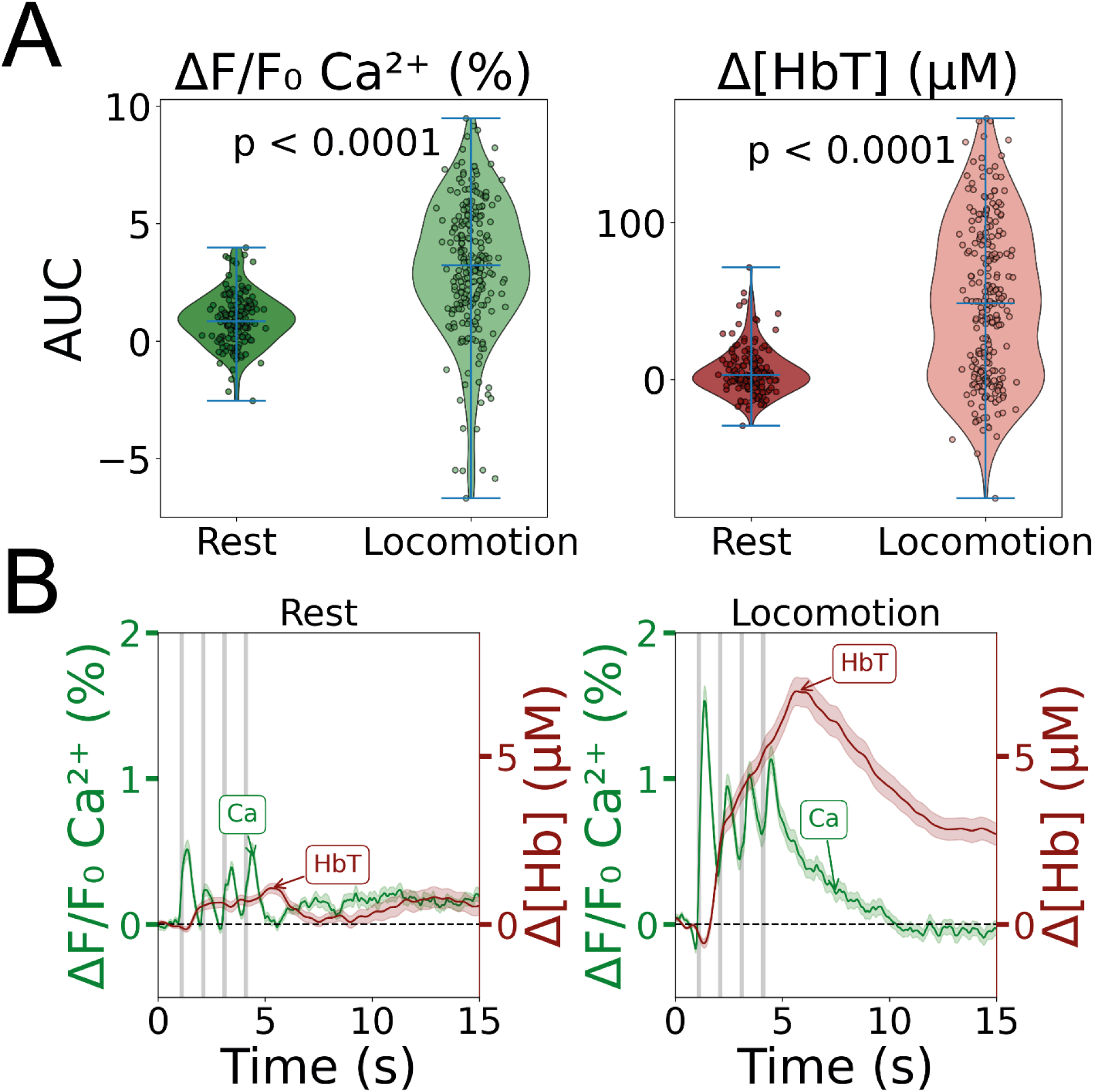
Behaviour significantly affects WFOI responses to stimulation. (**A**) Violin plots showing the area under the curve (AUC) of individual calcium and haemodynamic responses in the somatosensory cortex for hind limb stimulation during calm behaviour (Rest) and active locomotion (Locomotion). (**B**) Responses to hind limb stimulation: averaged Ca^2+^ signal from the response area (green line) and the corresponding haemodynamic response (dark red line), shown for both Rest and Locomotion conditions. Data were collected from 8 mice, with stimulation applied alternately to the left and right hind limbs in separate trials (21 repetitions of 4 pulses with 200 ms duration, 1 Hz frequency, I = 90 µA per limb); responses were recorded from the somatosensory cortex contralateral to the side of stimulation. P-values are based on Welch’s t-test.

These findings underscore the importance of monitoring behaviour during recordings. In this study, the behavioural state was assessed manually. However, deep learning-based tools, such as DeepLabCut (Mathis et al., 2018), could offer a more objective and reproducible approach for future behavioural classification.

## Discussion

There are two principal approaches to creating cranial windows: transcranial windows, which preserve the skull (Grinvald et al., 1986), and craniotomies, which involve skull removal (J. H. Kim et al., 1989). Two-photon microscopy typically requires either skull thinning (Kelly & Majewska, 2010) or removal with a glass-covered craniotomy (Holtmaat et al., 2009; Mikkelsen et al., 2022). WFOI, by contrast, benefits from the use of large cranial windows that expose up to 40–80% of the cortical surface.

While protocols for intrinsic imaging through the intact skull exist (Silasi et al., 2016; White et al., 2011), our observations suggest that variability in the diploic vessels may obscure neural signals. Additionally, the intact skull bone significantly reduces signal strength and the signal-to-noise ratio. Removing the trabecular (spongy) bone layer improves optical clarity and can be achieved via skull thinning (Shahsavarani et al., 2023) or skull replacement with transparent materials such as silicone (Manita et al., 2022), nanocoatings (Takahashi et al., 2021), PDMS (polydimethylsiloxane) (Heo et al., 2016; Yang et al., 2022), PET plastic (Ghanbari et al., 2019), polymethylpentene (Edelman et al., 2024), or glass (T. H. Kim et al., 2016). Chemical skull-clearing methods have also been developed (Cramer et al., 2021; C. Zhang et al., 2018).

Although skull-thinning techniques have been described (Kozberg et al., 2016; Shahsavarani et al., 2023), detailed protocols are often lacking. The method we present is technically simpler than full cranial replacement, carries a lower risk of infection, and minimises bleeding, when performed with proper skill. A key feature of our protocol is the use of a non-fluorescent, gel-based nail polish as a transparent and protective coating. While this approach is optimised for WFOI (particularly laser contrast imaging), it may also be compatible with two-photon microscopy. Potential limitations include the risk of neuroinflammation, dura mater opacification due to pressure, and long-term bone regrowth. The procedure is time-consuming at first but becomes efficient with practice. We demonstrated that the method is suitable for both recording evoked responses to sensory stimulation and chronic longitudinal imaging. The window remains optically stable and functional for extended periods, enabling repeated within-subject measurements.

Independent of the cranial window technique, stable head fixation is essential for high-quality imaging. This is typically achieved by attaching a fixation device during surgery, which is later mounted in a stereotaxic holder or a mounting frame under the microscope. Devices vary in form: screws, nuts (Silasi et al., 2016), plates (Kelly & Majewska, 2010; Manita et al., 2022; Takahashi et al., 2021), or custom clip-based holders (Ghanbari et al., 2019; Mikkelsen et al., 2022). We modified commercial headplates (Neurotar, Finland), originally designed to resemble helicopter blades, to increase cortical access while ensuring firm fixation. These lightweight, compact plates are compatible with the Mobile HomeCage® (Neurotar, Finland).

The WFOI system design varies depending on experimental goals. While commercial systems are available (e.g. Invigilo 3, Neurotar, Finland; OASIS Macro Cortex-Wide Intrinsic Optical Imaging, Mightex, CA, USA), many laboratories build custom setups that support intrinsic imaging, calcium, or FAD imaging, stroke induction, and other procedures (Shahsavarani et al., 2023; X. Wang et al., 2024; Zhao et al., 2021), including low-cost configurations (Harrison et al., 2009). Various solutions for the awake mice *in vivo* imaging exist: rotating disks (Heo et al., 2016; Nietz et al., 2022), spheres (Dombeck et al., 2007), cylinders (Cardin et al., 2020), treadmills (Fedotova et al., 2023; Moeini et al., 2020; Yi et al., 2024), hovercraft platforms (Juczewski et al., 2020; Stuart et al., 2024), and even hammocks (Padawer-Curry et al., 2023) or tubes (Silasi et al., 2016). WFOI is also compatible with virtual reality setups (Pinke et al., 2023).

We developed a flexible and cost-effective WFOI system using commercially available components, which can be readily adapted to various experimental needs. While high-end systems can exceed $200,000, our protocol outlines a minimal configuration suitable for constrained budgets.

We hope that this accessible and adaptable approach will promote the broader use of functional imaging techniques in neuroscience research, contributing to a deeper understanding of brain function and disease mechanisms.

## Supporting information

S2 SOPs

S3 3D models

S4 Troubleshooting

## Supplementary data

Supplementary data are available in Biology Methods and Protocols online.

## Data availability

Supplementary video materials are available on the YouTube channel “WIFIOPIA”: https://www.youtube.com/@WIFIOPIA.

The code described in this article and the sample data have been deposited in the GitHub https://github.com/natalia1896/WIFIOPIA). Any additional data, if needed, will be provided upon request.

## Author Contributions

**Kislukhina E.N**. (Methodology [equal], Validation [equal], Data curation [equal], Investigation [equal], 3D-Modelling [lead], Visualisation [lead], Writing-original draft [equal]); **Lizunova N.V**. (Methodology [equal], Validation [equal], Data curation [equal], Investigation [equal], Writing-original draft [supporting], Programming [lead]); **Surin A.M**. (Conceptualization [lead], Supervision [equal], Writing - review & editing [equal]); **Bakaeva Z.V**. (Conceptualization [equal], Data curation [equal], Supervision [lead], Writing - review & editing [equal], Project administration [lead]).

## Institutional Review Board Statement

The study was conducted in accordance with the ethical principles and regulatory documents recommended by the European Science Foundation (ESF) and the Declaration on Humane Treatment of Animals, and in accordance with the Order of the Ministry of Health and Social Development of the Russian Federarion no. 708n of 23 August 2010 “On Approval of Laboratory Practices”. Animal care, breeding, and experimental procedures were carried out as required by the Ethical committee of the Institute of General Pathology and Pathophysiology, Protocol no. 05-06/12 of 14 December 2017 and the Ethical committee of the National Medical Research Center for Children’s Health, Protocol no. 05 of 15 May 2025

## Acknowledgments

The authors gratefully acknowledge Prof. Piotr Brezhestovskiĭ for generously enabling access to the ClopHensor mouse line, and for his valuable methodological insights and suggestions. The authors thank Dr Leonard Khiroug, whose ideas and efforts provided the initial impetus for this research. Some portions of this manuscript, including language polishing, and draft code documentation (e.g., function docstrings), were assisted by ChatGPT (OpenAI, San Francisco, CA). Final decisions and edits were made by the authors.

