## Supplementary material for "The protocol for mesoscopic wide-field optical imaging in mice: from zero to hero": S2 SOPs

**Supplement materials**

### 1 Preparation of adaptor for headplate from epoxy resin

If the shape of the provided adaptor is unsuitable or if 3D-printing is unavailable, custom adapters can be made from epoxy resin.

1. Follow safety precautions: wear gloves and a respirator, and perform all procedures in a fume hood.
2. Make an impression of mouse skull with dental acrylic (e.g. Villacryl S, Zhermack, Poland). Apply dental resin around the intended cranial window layer by layer. For that, use a syringe with a 18G needle.

*It will be more convenient to use shortened 18G needle.*

3. Apply the last layer of the resin, and without allowing it to dry, position the flat metal blank headplate over it.
4. Wait for the resin to completely dry (about 10 minutes) and separate the headplate along with the resin adapter from the mouse's skull. File the existing roughness to achieve a smooth shape of master model.

*Additionally, the sides of master model can be coated with a layer of gel polish to enhance smoothness. Do not cover the underside, as it should remain slightly rough for secure adhesion to the scull.*

5. Make silicone molds from master model. Use a cropped plastic cup, cover its bottom with a thin layer of wax plasticine. Place the master model in the center of the bottom, flat side down. Press it lightly into the plasticine. Mix silicone components due to manufacturer's instructions (e.g. SF-20A-200, EpoxyMaster, Russia). Add 5 drops of the pigment (e.g. Epic Art Color, Epic Art, Russia). Pour silicone into the prepared plastic cups. Pour the silicone only onto the walls to prevent bubbling. Allow the silicone to cure, which usually takes 8 hours but may vary depending on the manufacturer and proportion of ingredients. Separate the plastic cup. Carefully remove the master model. You can reuse the master model to make more molds.
6. Mix epoxy resin (e.g. 476316047, ArtDealer, Russia). Using a syringe without a needle, measure the required amounts of both components of the epoxy resin in accordance with the manufacturer's instructions (2 ml of mixture per 20 headplate units). Add 1 drop of the black pigment (e.g. Epic Art Color, Epic Art, Russia) and mix them well in eppendorf tube.

*Take into account the shrinkage of epoxy resin. Due to the poor mixing of two-component polymers (silicone and epoxy resin) near the walls, after the initial mixing, the sample should be transferred to another container and mixed again. The addition of pigment is optional, but it helps to control the completeness of mixing. Try to minimize the presence of bubbles during the process*

7. Load the mixture into a clean syringe fitted with a modified (tip-removed) 18G needle. Fill the molds partially and insert metal headplate blanks, avoiding pressure that might introduce bubbles. Allow to cure fully (24–48 h) as per manufacturer's instructions, then remove the finished headplates.

#### 2 Wide-field thinned cranium surgery

**Preparation for surgery. Perform 1 day or more before surgery**

1. Cover all work surfaces, including equipment, with thick plastic wrap to prevent the entry of saline. Leave the tips of stereotaxis ear bars, the front of dental clamps, the lens of binoculars, the light source, and the tip of saline needles unwrapped.

*If any saline solution comes into contact with equipment, immediately dry the area and thoroughly rinse with distilled water to prevent rust formation.*

2. Create "skirts" for the milling cutters and boron. To accomplish this, cut out circles with a diameter of approximately 7 millimeters from transparent plastic using scissors, and create a hole in the center that corresponds to the diameter of the leg of the milling cutter. Place the "skirt" over the leg and secure it with gel polish approximately 5 millimeters from the bottom of the tip of the milling cutter.

*The gel polish should be applied on the side of the tip of the milling cutter, such that the saline solution can subsequently flow freely to the edges of the "skirt" and spread apart under the influence of the central force. If not, the solution will flow down the leg of the milling cutter into the drill mechanism.*

3. Dilute Marbofloxacin to a final concentration of 0.00017 mg/mL by adding 12 volumes of water for injection. Prepare aliquots in 1.5 mL Eppendorf tubes for future use.

*Marbofloxacin may precipitate in saline solutions. All solutions must be prepared aseptically and stored at +4°C.*

4. Dilute Ketonal by a factor of 30 with saline solution to achieve a concentration of 1.65 mg/ml. Prepare aliquots for future use.
5. Dilute Dexamethasone 11.4-fold with saline solution to achieve a concentration of 0.35 mg/ml. Prepare aliquots for future use.
6. Prepare angled cannula tips 18G, 45°. Using the needle holder, pinch the tip of the 18-gauge needle and swing it from side to side to break off the tip. The remaining length of the needle should be approximately 1.5 centimeters. Using a needle holder, bend the distal 4 millimeters of the needle by 30 degrees without pinching the lumen.
7. Prepare a 27-gauge L-shaped needle for dispensing glue and removing bubbles from the polish. To do this, use a needle holder to bend the needle 90 degrees along its length, and then bend the tip an additional 90 degrees (toward the base) to avoid self-injury in the future when handling it. Place the needle on a syringe or any suitable long stick.
8. Prepare semi-C-shape 27G needle for applying black lacquer markings on the surface of cranial window by breaking off the tip of a 27-gauge needle and bending it into an arc.
9. Cut the gauze into strips approximately 2 x 6 cm for drainage purposes.
10. Provide a supply of cotton swabs/gauze to clean the binocular lenses.
11. Make visor from aluminum foil to protect mouse's eyes from UV light.

*It would be convenient to keep the listed small supplies in one place.*

12. Extend a silicone tube from the peristaltic pump to the stereotactic device. Attach a blunt medical needle to the distal end of the tube. Secure the needle to a wire or other three-axis-movable device, allowing for fine adjustment of its position during surgery.
13. Extend the silicone tube from the air pump to the stereotaxis. Fill the tip from the 1000 µl pipette with filter material (cotton swab). Attach the tip to the tube.

**Preparation for surgery. Perform on the day of the operation**

14. Check the availability of all necessary tools and medication.

15. Disinfect the workplace with 70% ethanol.

16. Disinfect tools and screws.

*Headplates with epoxy resin adapters should not be sterilized using high temperatures or alcohol, as the resin will become soft. Instead, they can be treated with 70% ethanol prior to gluing.*

17. Prepare a reservoir with distilled water, a nail stick, and cotton swab to wash off the depilatory cream.

18. Prepare a clean cage with fresh water and food.

19. Rinse the perfusion pump tubing systems (you may turn the pump up to maximum speed):
    a. Peroxide (50 mL).
    b. Distilled water (50 mL).
    c. Fill it with sterile saline.

*Rinsing is not necessary if several procedures are performed in succession.*

20. Setting up the perfusion pump:

a. Hang the 200 mL saline solution package, lid down. It is important to ensure that there is no risk of leakage of saline solution onto the equipment.
b. Connect the 18-gauge needle base to the tubing of the perfusion pump and insert the needle into the saline solution container.
c. Set the flow rate to 1–2 mL/min (while the pump is off). *This can be adjusted to individual preferences.*

21. Turn on temperature-controlled stage for animals (37 °C).

**Animal preparation**

22. Anesthetize the animal:

a. Ensure that the pumps of the isoflurane system are turned on. Verify that the $O_2$/air flow to the anesthesia mask is 1 liter per minute, both before and after passing through the vaporizer.
b. Connect the supply line to the induction chamber and set the concentration of isoflurane at 4%.
c. Place the mouse into the induction chamber for approximately 3 minutes.
*During this time, monitor the animal's breathing rate. When the breathing rate decreases to approximately 1 breath per second, remove the mouse from the chamber. If breathing becomes convulsive or intermittent, remove the animal immediately from the chamber to prevent further complications.*

23. Gently retracting the vibrissae with one finger, shave the fur on the dorsal surface of the head using a razor.

*If the mouse begins to awaken after shaving, it can be placed back into the induction chamber.*
*A thorough shave is not necessary, removing the majority of the fur from the headplate site is sufficient.*

24. Convert isoflurane to 2%.

*Depending on the line and the individual characteristics of the animal, it is necessary to choose 1-2.5% isoflurane to maintain anesthesia. It is necessary to maintain regular breathing for about 1 breath / s and control the absence of a pedal reflex.*

25. Secure the mouse within the stereotaxic device.

*Using forceps, fix the teeth in the holder's opening, slide the mask in place, and secure it. Use the ring or little fingers to move the stereotaxic clamps, being careful not to damage the mouse's skull. Ensure that the clamps rest against the temporal depressions anterior to the ear openings.*

26. Apply eye gel to the mouse's eyes, repeating as necessary to prevent drying (approximately every 15 minutes if saline administration is not used).
27. Administer intramuscular dexamethasone (dose: 0.7 mg/kg; volume: 2 μL per gram body weight).
28. Lubricate the rectal probe and insert it into the anus of the mouse, securing it with tape around the tail.
29. Subcutaneously inject 0.1 mL of 0.2% lidocaine under the scalp of the mouse.
30. Place cotton beneath and around the mouse's head to absorb any leaking saline solution.
31. Adjust the microscope and lighting for binocular viewing.

**Preparation of the surgical field**

32. Apply depilatory cream to the scalp for 1 minute, covering an area 1–2 mm wider than the planned incision.

*If the cream is not fresh, extend the application time to 5 minutes. Minor skin damage is not critical, as the skin will be removed during the procedure.*

33. Remove the cream and fur by scraping them off with the flat end of a wooden manicure stick. Wipe away any remaining residue with a cotton swab, then clean the skin with a water-soaked cotton pad.
34. Disinfect gloves with 70% ethanol.
35. Disinfect the mouse’s head by wiping it three times with a 70% ethanol-soaked cotton swab.

*Ensure that the ethanol does not come into contact with the eyes.*

36. Confirm the absence of a pedal reflex.
37. Excise the scalp using scissors and forceps, ensuring smooth, non-torn edges. The incision should expose half of the occipital bone, the parietal and frontal bones up to the muscle attachment sites, and approximately 1 mm of the nasal bones (up to the anterior eye line).

*The skin will naturally stretch over the occipital region.*

38. To control capillary bleeding, apply haemostatic agent to the wound edges and allow it to dry.
39. Remove the fascia using a cotton swab or scalpel.

*It must be completely cleared from the exposed area, ensuring that no remnants are trapped beneath the implant headplate, as this would compromise fixation. The edges should also be entirely removed to prevent them from becoming entangled in the burr drill during subsequent procedures.*

**Thinning of the skull**

40. Place a piece of gauze (drainage) over the mouse’s nose so that its edges are in contact with the cotton, while the central part rests on the nasal bones. Use the same gauze to cover and retract the vibrissae downward, as well as to protect the eyes.

*The central portion of the gauze should be in direct contact with the surface of the nasal bones and positioned no closer than the external cerebral vein (the boundary between the nasal and frontal bones). For convenience, pre-soak the gauze in saline before placement.*

41. Ensure a stable fixation of the mouse's head in the stereotactic device. If necessary, further secure the ear bars.

*A more secure head fixation reduces the risk of brain damage.*

42. Turn on the peristaltic pump at a flow rate of 1–2 mL/min and check that the drainage effectively removes excess fluid.

*Without proper drainage, saline may flow down the fur into the mouse's mouth and nose, increasing the risk of aspiration.*

43. Adjust the position of the peristaltic pump's needle so that saline droplets fall rostral to the area being thinned.

*If the droplet falls onto the cutter or caudally, the cutter may disperse the liquid, contaminating the binocular microscope's lenses.*

44. Set the microdrill speed to approximately 25,000 rpm.

*The speed can be adjusted based on the surgeon's preference and the mouse's characteristics (younger mice require lower speeds). The speed may also be reduced when working with the red (soft) dental cutter.*

45. Insert the diamond cutter into the microdrill.

46. Rest two fingers of your non-dominant hand on the stereotaxic frame on both sides of the mouse while pulling back its ears. Place the edge of your dominant hand, which holds the drill, on the heating pad, stereotaxic frame, or your other hand.

*This helps stabilize your hand position and prevents damage to the mouse's ears.*

47. Turn on the microdrill.

*Foot pedal-operated models are more convenient.*

48. Use the tip of the dental cutter to grind the frontal bones along the perimeter. Bleeding from diploic vessels may occur at this stage; move to a different area and allow it to stop before proceeding.

*The resulting bleeding comes from diploic and emissary vessels, which should be removed during thinning.*

*However, avoid bleeding from subcortical vessels located beneath the bone tissue. If excessive bleeding occurs, dry the skull with an air stream and apply a Hemostab-soaked cotton swab.*

49. Continue thinning the entire surface of the frontal bones. Maintain tangential motion for uniform thinning.

*The drill angle determines the uniformity of thinning, which is critical for a successful procedure.*

50. Slightly thin the sutures, but avoid full thinning of the anterior ¼ of the outer sagittal sinus, as this may cause severe bleeding. Minimal thinning should be performed in this area.

51. Turn off the microdrill and peristaltic pump. Turn on the air pump and dry the skull with an air stream. Visually assess the uniformity and degree of skull thinning.

*A fully thinned skull appears pinkish when dry. Unthinned areas remain white. When thinning the frontal bones, the parietal areas are often affected as well.*

*Uneven thinning (where pinkish thinned regions border white unthinned areas) increases the risk of skull fractures. To prevent this, use continuous circular or linear motions and avoid contacting bones other than the targeted area with the lower part of the milling cutter. If a crack forms, change the drill's rotation direction and avoid touching the damaged site.*

52. Turn on the peristaltic pump and thin the parietal bones. Thinning can be performed using continuous movements from temple to temple along the sagittal suture. Periodically dry the skull to assess the degree of thinning.

53. Further refine the sutures.

54. Using the diamond milter cutter, followed by a medium (blue) silicone-diamond cutter, remove the outer compact bone layer and most of the spongy bone, leaving the inner compact layer intact Once the skull appears pinkish upon drying, switch to a soft (red) dental milter cutter and perform polishing movements over the entire skull surface.

55. Switch to an extra-soft (gray) dental milter cutter and further polish the entire skull surface.

*The diamond and blue* dental milter cutters *are effective for thinning the sutures. The red and gray* cutters *are used for polishing.*

56. Continue thinning until the dry and wet skull appear equally transparent, with clearly visible blood vessels.

*A fully thinned skull should not be left dry for extended periods.*

57. Use scalpel blade №15 to score nasal bones for proper headplate fixation.

**Application of Coating and Headplate Fixation**

58. Turn off the microdrill and peristaltic pump. Reapply eye gel to the mouse's eyes.

59. Dry the skull with an air stream. Apply a small drop of cyanoacrylate glue to an L-shaped bent needle and carefully distribute a thin layer only over the thinned surface. Allow the glue to set.

*To speed up drying, an air stream can be used.*

60. Try to fit the headplate. Confirm head stability. If necessary, tighten the stereotaxic screws further.

61. Drill a 1-mm in diameter, ~0.3 mm deep indentation in the occipital bone caudal to the headplate border (in the lower central portion of the occipital bone). Use tungsten carbide drill bit.

62. Insert a M1 screw into a pre-drilled indentation. Use screwdriver and curved forceps to fix it.

*Bones are soft enough to allow screw thread to penetrate it. Continue tightening only as much as necessary to prevent wobbling (it will not be possible to secure it very firmly). Do not penetrate the bone completely, as this may trigger cerebellar inflammation.*

63. Sterilize the headplate with 70% ethanol and test its fit on the skull.

64. Apply glue to the inner rim of the headplate. Gently press the headplate onto the skull and allow the glue to set for 10–15 seconds.

*Apply light pressure only, using the weight of the hands rather than muscle force, to prevent skull damage.*

65. Place an aluminum foil shield over the eyes to protect them from UV exposure.

66. Apply a base coat of gel polish to the entire thinned surface using a brush or an L-shaped needle.

*Avoid covering the area where the headplate will be attached. The layer should be as thin as possible and free of bubbles.*

67. Cure the base coat under a UV lamp for 10 seconds, then pause for 10 seconds, followed by 60 seconds of curing.

*Since the gel polish generates heat during polymerization, avoid thermal damage to the brain.*

68. Apply a top coat of gel polish over the base coat and cure under the UV lamp for 10 seconds, followed by up to 60 seconds.

69. Using a curved blunt needle, mark the bregma with black nail polish.

*Proper thinning renders the bregma visually indistinct.*

70. Prepare the acrylic resin. Use a spatula to scoop about 0.25 g of powder into a silicone mortar. Using an insulin syringe (without the needle), add 200-250 µL of solvent. Mix thoroughly and load into a syringe fitted with an 18G needle, with the tip broken off and bent at a 45° angle.

71. Apply the acrylic resin one drop at a time along the outer perimeter of the headplate, allowing each drop to set.

*This prevents the resin from spreading over the cranial window.*

72. Seal the entire perimeter of the headplate with acrylic resin, covering the skin edges. Ensure a thick layer over the nasal and occipital bones. Do not touch the mouse's eyes with acrylic resin.

*Avoid excessive resin on the sides, as this may prevent proper mounting in the holder.*
*Bleeding may resume. Try fully stop the bleeding, as blood can compromise the acrylic's solidity.*
*If resin or nail polish spreads over the surface of the cranial window too much, allow it to partially dry and remove excess using a needle or a small acetone-soaked cotton swab.*

73. After the resin hardens, remove the mouse from the stereotaxic apparatus. Subcutaneously administer Antibiotic (Marbofloxacin, D = 8 mg/kg, V = 150 µL/30 g mouse), analgesic (Ketoprofen, D = 8.5 mg/kg, V = 150 µL/30 g mouse), 0.5 mL of saline solution.

*Marbofloxacin precipitates in saline, so administer it using a separate syringe.*
*Administer injections near the tail to avoid tension and trauma to the head skin.*

74. Turn off isoflurane delivery and place the mouse in a clean, heated single-housing cage Monitor the animal closely for at least one-hour post-surgery.

75. For the first three days post-surgery, subcutaneously administer Marbofloxacin and Ketonal once daily while monitoring and documenting the animal’s condition, check for signs of inflammation.

*Well-handled mice should not display aggressive behavior during injections*
*The rehabilitation period should last at least 14 days, though a one-month recovery is recommended for full restoration.*

#### 3 Mouse habituation to head fixation

1. Turn on the setup. The noise level in the room during habituation should match that of the actual experiment.
2. Transfer the cage with the mouse to the experimental room and allow the animal to acclimate for at least 30 minutes.
3. Handle the mouse and allow it to calm down.

*Note: Mice should be well-acclimated to handling before.*

4. Gently restrain the head for 5 seconds. Repeat twice.

*If the mouse resists strongly, release it earlier.*

5. Gently restrain the head for 10 seconds. Repeat thrice.
6. Transfer the mouse to the setup and allow it to explore the movable cage for 1 minute.

*Prevent escape attempts using your hand.*

7. Return the mouse to the home cage.
8. On the following day, repeat steps 1–5, skipping step 4.
9. Adjust the airflow so that a sheet of paper can be slid under the cage.
10. Then gently restrain the mouse's head and bring it close to the head-fixation apparatus. Hold the mouse near the apparatus for 5-10 seconds. Repeat three times.
11. Return the mouse to the home cage.
12. On the following day, turn on the setup, bring the mouse into the room, and allow 30 minutes for acclimation. Handle the mouse, allow it to calm down, transfer it to the setup, and let it explore for 30 seconds. Gently restrain the head and bring the mouse close to the fixation apparatus for 5-10 seconds.
13. Secure the mouse by the head cap and fix it in the head holder. Tighten the screws and release the mouse after 10 seconds. Repeat three times.

*Adjust the height of the fixation apparatus so that the mouse's head is fixed at a height close to its natural posture.*

14. Fix the mouse in the apparatus and lower the camera to the imaging position.
15. Clean the skull with a lint-free tissue moistened with 70% ethanol, cover the eyes with a visor, and turn off the room lights. You can turn on LEDs to imitate the condition of the experiment.

*Skull cleaning is not required if protective caps are used.*

16. Wait for 60 seconds.
17. Turn on the light and return the mouse to the home cage.
18. On the following day, record the resting-state activity for no more than 5 minutes, as described below, and return the mouse to the home cage.

*After this habituation protocol, the mouse should remain calm during head fixation and should not associate the setup with aversive stimuli.*

#### 4 Imaging procedure

1. Power on all devices.
2. Turn on the computer.
3. Transfer the mouse to the experimental room and allow 30 minutes for acclimation.
4. Load the imaging protocol and camera settings. Set the file-saving path and enable automatic file naming with time stamp.

*You can use automatization, such as AutoHotkey v2.0 software (Indiania, USA) to avoid human inaccuracy.*

5. If working distance of your lens is small, raise the camera.
6. If necessary, induce anesthesia using 4% isoflurane and turn on heating pad.
7. Position the mouse in the setup and, if necessary, administer 1.5% isoflurane via a mask.
8. Secure all screws with a screwdriver.

*Hand-tightening may be insufficient.*

9. Clean the skull with a lint-free tissue moistened with 70% ethanol (or remove the protective cap if used).
10. Cover the mouse's eyes with the visor.
11. Set the camera to continuous mode, preview images, and activate an LED in continuous mode.
12. Lower the camera and focus it on the center of the hemispheres.

*It is not possible to perform equal focus for whole hemispheres due to their form.*

13. Turn off the room lights.
14. Adjust the camera exposure to avoid glares formation. Write down 70–80% of the exposure time at which glare first appears.

*If our 3D-printed mount for 1-7 optic bundle is used, some holes should be covered with diffusor (e.g. white paper medical tape) to prevent glare.*

15. Repeat step 14 for all LEDs.
16. Restore the initially selected camera exposure.
17. Adjust LED pulse durations based on the defined values (step 14).
18. Switch the camera and LEDs to trigger mode.
19. Start imaging.

*Remain behind the mouse during imaging, avoid movement, noise, or changes in lighting, and prevent vibrations. Any sensory stimulus will affect brain activity.*

20. If necessary, record multiple sessions.

*Total head-fixation duration should not exceed 2 hours, with 1 hour being optimal.*

21. After imaging, turn on the lights, release the mouse, and return it to the home cage.
22. Transfer raw data to a secure storage drive.
