## Supplementary figures and images for "The protocol for mesoscopic wide-field optical imaging in mice: from zero to hero"

### Headplate.tif

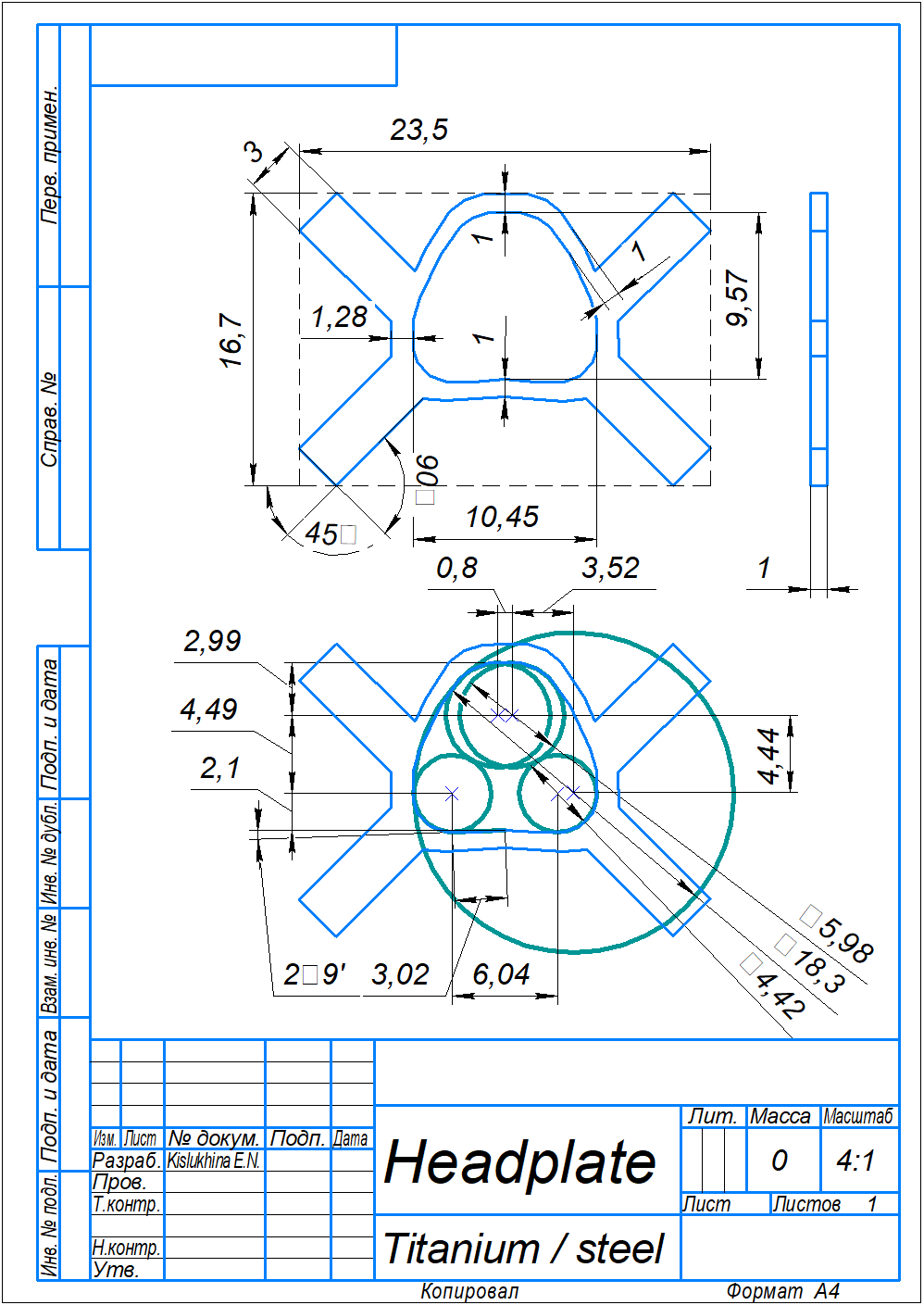
