## Supplementary material for "The protocol for mesoscopic wide-field optical imaging in mice: from zero to hero": S4 Troubleshooting

**Troubleshooting guide for wide-field cranial window surgery and wide-field optical imaging**

| Problem | Possible Cause | Solution |
|---|---|---|
| Bone overgrowth | Incomplete thinning of the growth zone near sutures | Carefully thin the sutures and borders of the frontal bones, polish the sutures |
| Bone cracks | Uneven thinning | Ensure uniform and gradual thinning of all bones in the window area during surgery. Use circular or linear drill movement trough large area. Stabilise working hand |
| Bubbles in coating | Excess gel polish that did not distribute evenly, overly fast brush movements causing foaming | Apply a thin layer of polish, remove excess of polish from brush prior to covering |
| Bleeding during surgery | Vessel damage near sutures, bone cracks | Minor bleeding usually stops on its own; for severe bleeding, use capillary hemostatic agents or drop of cyanoacrylate glue |
| Eye damage | Contact with irritating substances (alcohol, solvent), insufficient eye hydration during surgery, excessively bright light | Use ophthalmic gel for eye hydration; reapply as needed. Be careful when applying dental acrylic or other irritants near the eyes. Use foil visor for surgery, 3D-printed visor for imaging |
| Intense wound scratching | Inflammatory response in the wound area | Use anti-inflammatory agents. To minimize scratching of unhealed wounds, consider a veterinary collar for no more than 3 days post-surgery |
| Unstable LED brightness | Cross-talk between channels of multi-channel controller. Unstable current supply. LED damage | Set 50% or more of LED intensity (mount ND filters if needed). Use isolated stimulator. Use proper UPS. |
| Hemispheres exposed to external light | Insufficient contact between the adapter and the skull, and inadequate light shielding between the stimulating eye LED and the cranial window surface | Check the quality of the adapter. If any openings are present, cover them with dark nail polish. Add an additional light shield if necessary. |
| Incorrect number of saved frames | Improper camera settings, insufficient memory or processor processing power, unstable cable connections (damaged ports) | Identify the issue, check cable connections, restart the computer |
| Headplate detachment during experiment/training | Loose fixation during surgery, inflammation, excessive force from the experimenter/mouse | If skull damage is minimal and the experimental protocol permits, reattach the headplate (preferably with a screw) and ensure proper postoperative recovery |
